# Characterization of both major histocompatibility complex classes in a wild social mammal: the banded mongoose

**DOI:** 10.1101/2024.12.14.628504

**Authors:** Nadine Schubert, Hazel J. Nichols, Francis Mwanguhya, Robert Businge, Solomon Kyambulima, Kenneth Mwesige, Michael A. Cant, Jamie C. Winternitz

## Abstract

The major histocompatibility complex’s (MHC) role in the vertebrate adaptive immune response and its exceptional polymorphism make it a key target for studying adaptive gene evolution. However, previous studies on carnivore MHC have mostly focused on populations which experienced a severe bottleneck or are of general conservation concern. Hence, sample sizes are often small and generalizations about MHC diversity are unreliable. Furthermore, studies often focus on one MHC class and do not cover the whole peptide binding groove of the MHC molecule. Here, we characterize MHC class I (MHC-I) exon 2 and 3, encoding both the α1- and α2-domain of the MHC-I molecule, as well as MHC-II DRB exon 2 for a large sample (N = 282-485) of a wild mammal of least conservation concern, the banded mongoose. We found that MHC-I generally showed higher allelic diversity and polymorphism compared to MHC-II, which is in line with findings in humans that show higher diversifying selection acting on MHC-I. However, MHC-I exon 3 showed the lowest diversity, possibly due to its different role in generating the peptide binding groove of the class I molecule compared to exon 2. Moreover, we found selection to act more strongly on MHC-I exon 2 (domain α1) than exon 3 (domain α2). Despite frequent inbreeding, phylogenetic comparative analysis showed banded mongooses to have MHC diversity levels comparable with other carnivores of least concern. Phylogenetic analysis indicated a longer evolutionary trajectory for MHC-II compared to MHC-I as well as species-specific gene duplication of nonclassical sequences of MHC-I clustering with classical sequences. Trans-species polymorphism was detected for nonclassical MHC-I sequences suggesting homology or convergent evolution for these genes. Our study is the first to characterize both MHC classes of a social, wild carnivore using a high throughput sequencing approach with a large sample size and thereby provides the basis for further investigation of MHC structure and function within the banded mongoose and other carnivores.

## BACKGROUND

Due to their role in the immune response towards pathogens (Piertney and Oliver 2006; Bitarello et al. 2018; Ebert and Fields 2020), immune genes exhibit an exceptionally high level of polymorphism in both vertebrates and invertebrates (Ebert and Fields 2020). In jawed vertebrates the major histocompatibility complex (MHC) plays a crucial role in the adaptive immune response (Kaufman 2018). It encodes membrane glycoproteins that bind and present peptides and thereby facilitate discrimination between self and foreign peptides (Bjorkman et al. 1987; Knapp 2005).

MHC molecules are divided into two classes. Class I molecules (MHC-I), which are expressed on almost all nucleated cells, mainly present peptides found in the cytoplasm to cytotoxic T-cells. After activation, these T-cells induce the death of the MHC-I-carrying cell (Klein 1986). In contrast, MHC-II molecules are only expressed by professional antigen-presenting cells (APC), such as B-cells, macrophages and dendritic cells, that present peptides that have been ingested by the APC (Neefjes et al. 2011). This causes MHC-I to mostly bind the organism’s own self-peptides or antigens stemming from pathogens such as viruses that have entered the cell whereas MHC-II predominantly presents peptides of extracellular origin, such as antigens originating from parasites, that have been engulfed by the APC (Rammensee et al. 2013).

Within the two MHC classes, MHC molecules can be further subdivided into classical and nonclassical molecules. Classical MHC molecules have three key features: they are highly expressed, highly polymorphic, and specialized in presenting peptides to T-cells (Braud et al. 1999; Alfonso and Karlsson 2000). Nonclassical MHC molecules, while structurally similar to classical ones, have diverse functions. These include antigen processing, immunomodulation in both innate and adaptive immunity, and even roles unrelated to immune responses (Braud et al. 1999; Alfonso and Karlsson 2000; Adams and Luoma 2013). Given our interest in the relationship between MHC molecules and pathogen response, we focus on classical MHC molecules, which play a central role in immune defense.

A functionally crucial part of MHC molecules is the peptide binding region (PBR) which includes the amino acid residues responsible for binding antigens. In both classes the PBR is a dimer built by two domains: MHC-I PBR is formed by α1 and α2 domains (encoded by MHC-I exon 2 and 3) (Saper et al. 1991) whereas MHC-II is a heterodimer consisting of an α1 and a β1 domain (encoded by MHC-II genes A and B) (Brown et al. 1993). We focus on the DR β1 domain because the DR α1 domain is functionally monomorphic in carnivores (Yuhki and O’Brien 1997).

The peptide-binding region (PBR) of MHC molecules plays a crucial role in binding antigens and triggering immune responses. This central function subjects the PBR to high rates of amino acid substitutions, resulting in significant sequence polymorphism (Saper et al. 1991; Brown et al. 1993). MHC molecules with different PBRs have varying affinities for pathogenic peptides, so carrying a diverse range of MHC alleles encoding various MHC molecules is thought to be advantageous (Pierini and Lenz 2018).

This idea has led to two key theories: “heterozygote advantage hypothesis” (Hughes and Nei 1989; Takahata and Nei 1990) and the “rare allele advantage hypothesis” (Clarke and Kirby 1966; Zinkernagel and Doherty 1975). In the first hypothesis, heterozygous individuals, with more diverse MHC alleles, can respond to a broader range of pathogens more effectively than homozygous individuals. In the second hypothesis, pathogens adapt to target common alleles, making rare alleles more beneficial for their carriers. This hypothesis considers the dynamic nature of pathogen pressures over time.

Another mechanism that contributes to MHC polymorphism is birth-and-death evolution. In this process, new genes are generated through duplication. Some remain functional and persist in the gene pool for long periods, while others accumulate harmful mutations and become non-functional pseudogenes (Nei et al. 1997).

Pathogen-mediated balancing selection acting on the MHC is believed to be the main mechanism responsible for population allelic diversity (Bergström and Gyllensten 1995; Radwan et al. 2020) as well as trans-species polymorphism, where alleles are conserved and shared across speciation events (Klein 1987; Těšický and Vinkler 2015). This maintenance of diversity of the MHC has been observed even in cases where overall genetic diversity is extremely low, emphasizing its outstanding key role in determining individual fitness (Aguilar et al. 2004).

The MHC plays a key role in the immune system, and its diversity is closely linked to fitness. Understanding MHC diversity, polymorphism, and selection is especially important for species that have faced severe population bottlenecks (e.g. Castro-Prieto et al. 2011) or are at risk of extinction due to very small population sizes (e.g. Aguilar et al. 2004). In general, large, natural populations are expected to maintain greater MHC diversity and heterozygosity compared to populations that have experienced bottlenecks (Klein 1986; Hedrick 2003). However, even populations that went through severe bottlenecks may retain significant functional diversity in their MHC alleles (e.g. Gutierrez-Espeleta et al. 2001; reviewed in Radwan et al. 2010). This can occur because MHC molecules can bind a wide and overlapping range of peptides, masking a loss of diversity (Rao et al. 2011). Therefore, studies should not rely solely on measuring allelic diversity but should also use methods that assess functional diversity, such as identifying MHC supertypes (e.g. Reche and Reinherz 2007).

The banded mongoose (*Mungos mungo*) is a wild carnivore classified as being of least conservation concern. However, the recent emergence of *Mycobacterium tuberculosis* complex infections in this species―particularly *M. mungi* (Alexander et al. 2015; Alexander et al. 2016; Alexander et al. 2018) and *M. bovis* (Brüns et al. 2017)―highlights the need for a better understanding of its immune system. Furthermore, the banded mongoose has an unusual social and mating system which results in frequent inbreeding. Banded mongooses are cooperative breeders that live in social groups of 10-40 adults (Cant et al. 2016). In contrast to most cooperative breeders, there is low reproductive skew and members of both sexes reproduce regularly. Unusually, breeding is synchronized within each social group, and multiple females give birth together, usually on the same night (Hodge et al. 2011). The resultant pups are combined into a single litter and are raised communally by the group, but there is no evidence that mothers are able to recognize their own pups or vice versa (Marshall et al. 2021). In the best studied population of banded mongooses (in Queen Elizabeth National Park, Uganda), individuals show limited dispersal and often stay and breed within their natal group, resulting in close relatives often being among the pool of potential mates (Nichols et al. 2014). As the mongooses’ ability to recognize relatives behaviorally is limited (Khera et al. 2021), inbreeding is common. For example, a pedigree of the Ugandan population showed that the majority of individuals (66.4%) were to some extent inbred, with 7.1% being closely inbred (resulting from full sibling or parent-offspring matings) (Wells et al. 2018), which results in substantial inbreeding depression (Wells et al. 2018; Wells et al. 2020). However, banded mongooses may reduce inbreeding by preferentially mating with individuals that are less closely related to them than expected from random mate choice (Sanderson et al. 2015). Moreover, they seek out extra-group mating opportunities, which can reduce their inbreeding levels (Nichols et al. 2015), but mechanisms of mate choice remain unclear (Khera et al. 2021). A long-term study on banded mongooses provides this unique opportunity to investigate MHC diversity, polymorphism and selection patterns in a social wild mammal with high potential for inbreeding.

Using next generation sequencing (NGS) with a stringent protocol for high throughput sequencing, we aim to gain a better understanding of MHC evolution in carnivores. Specifically, we (i) characterize the PBR for both classes by sequencing MHC-I exon 2 and 3, and MHC-II DRB exon 2 and determine their levels of polymorphism, (ii) compare MHC diversity and polymorphism with that found in other carnivores while controlling for phylogeny, (iii) identify patterns of selection acting upon MHC genes, and (iv) conduct phylogenetic analysis among related carnivores to determine potential trans-species polymorphism.

## METHODS

### 2.1 Study site and population sampling

Data were collected between 1994 and 2018 from a wild population of banded mongooses inhabiting Queen Elizabeth National Park in Uganda (0°12’S, 27°54E’). The study area covered approximately 10 km² savannah including the Mweya peninsula and the surrounding area. The population contained approximately 10-12 social groups, corresponding to approximately 250 individuals alive at one point. Banded mongooses were captured in Tomahawk traps (Tomahawk Live Trap Co., Tomahawk, WI) and were anesthetized with isoflurane as described in detail Jordan et al. (2010). A 2-mm tissue sample for genetic analysis was taken from the tip of the tail using sterile surgical scissors. A diluted solution of potassium permanganate was then applied to the wound to minimize risk of infection. Tissue samples were stored in 96% alcohol until further processing. All animals were given a tattoo or a subcutaneous pit tag (TAG-P-122IJ, Wyre Micro Design Ltd., UK) to allow permanent identification. Animals were then allowed to recover in a covered trap with access to water. Following this, they were released together with other members of their social group at the site of capture.

### 2.2 Ethical statement

Research was conducted under approval of the Uganda National Council for Science and Technology, and all procedures were approved by the Uganda Wildlife Authority and the Ethical Review Committee of the University of Exeter. All research procedures adhered to the ASAB Guidelines for the Treatment of Animals in Behavioural Research and Teaching (ASAB Ethical Committee and ABS Animal Care Committee 2022).

### 2.3 Genetic analyses

#### 2.3.1 DNA extraction

We extracted DNA from 465 individuals using Qiagen® DNeasy blood and tissue kits (Quiagen, the Netherlands) according to the manufacturers protocol. DNA was stored in the provided elution buffer or TE buffer and kept frozen at -20 °C until further processing.

#### 2.3.2 MHC genotyping

##### 2.3.2.1 Amplification of MHC genes with high-throughput sequencing

The extracted DNA was used to amplify fragments of MHC-I exon 2, MHC-I exon 3, and MHC-II DRB exon 2 of banded mongooses. Primers for MHC-I exon 2 were established based on the felid sequences published by Yukhi & O’Brien (1990), whereas the primers of MHC-I exon 3 and MHC-II DRB exon 2 were newly designed based on the published sequences of other closely related carnivore species (NCBI accession numbers can be retrieved from Table S1 in the supplementary material). We used these primers to amplify the 271-bp fragment (228-bp excluding primers) of MHC-I exon 2 (forward: CCACTCCCTGAGGTATTTCTACACC, reverse: CTCACCGGCCTCGCTCTG), the 270-bp fragment (232-bp excluding primers) of MHC-I exon 3 (forward: GGTCACACAGCATCCAGAGA, reverse: GCTGCAGCGTCTCCTTCC), and the 239-bp fragment (201-bp excluding primers) of MHC-II DRB exon 2 (forward: CGAGTGCCATTTCACCAACG, reverse: GCTGCACCGTGAAGCTCT). Furthermore, we added 10 bp tag sequences with an edit distance of 7 to the primers, so that a combination of 8F- and 12F-tags would enable unique combinations for each sample (Faircloth and Glenn 2012) (Table S2). This way we were able to pool the amplicons of the samples of a plate after the PCR and use one Illumina adapter per pool while still allowing unique identification during de-multiplexing, clean-up and processing. In addition, primers included 6-8 bp ‘junk’ sequences added to the 5’ direction (Table S2) to reduce inference during sequencing-by-synthesis through strong signals caused by a high number of the same bases being sequenced simultaneously. The complete primer consisted of the ‘junk’ sequence followed by the tag sequence with a specific forward or reverse primer sequence for one of the MHC fragments (5’-junk-tag-primer-3’).

We amplified each sample in duplicate and ran a negative control in a differently positioned well within each plate to avoid potential amplification bias caused by the differing primer pair combinations. PCRs were carried out with a total reaction volume of 25µl containing 14.3 µl of DNase and RNase free water, 2 µl of Buffer S (VWR, United States), 0.5 µl 40 mM dNTPs (Carl Roth, Germany), 0.5 µl of the tagged forward and reverse primers diluted 1:10, and 2 µl of DNA (corresponding to ∼40-100 ng of DNA). The PCRs were run on cyclers according to the following protocol: denaturation at 94°C for 2 min, followed by 30 cycles of 94°C for 30 seconds, annealing at either 55°C for MHC-I exon 2 and 3 or 58°C for MHC-II DRB exon 2 for 30 seconds, and elongation at 72°C for 60 seconds, and finally 10 min of elongation at 72°C and cooling down to a storage temperature of 4°C. We monitored the success of the PCR by running 5 µl of the PCR product on a 2% agarose gel.

We used the NGS Normalization 96-well kit (Norgen Biotek Corp., Canada) according to the manufacturers protocol to clean up the PCR products and simultaneously adjust amplicon concentrations. We then pooled the normalized samples per plate for preparation of libraries and measured the amplicon concentrations of the pools using the Qubit® dsDNA High Sensitivity Assay Kit on a Qubit® 2.0 Fluorometer (Invitrogen/Thermo Fisher, United States). We used the Illumina TruSeq DNA Nano Low Throughput Library Prep Kit (Illumina Inc, United States) in combination with the TruSeq DNA Single Indexes Set A and B (Illumina Inc, United States). Some indexes were used for multiple libraries, but together with the unique combination of tags and primer sequences, we ensured identification of samples during high-throughput sequencing. We followed the manufacturers protocol for library preparation with some minor adjustments. Due to lower concentrations of the pooled amplicon samples (ranging from 0.04 ng/µl to 0.3 ng/µl) we increased the amplicon volume in the end repair step to 200 µl instead of 50 µl. Furthermore, we prepared dilutions of the indexes used for each pool according to the amplicon concentration with an undiluted index volume being adjusted to 100 ng of DNA to avoid dimers.

We checked library quality using the Agilent Bioanalyzer 2100 (Agilent Technologies, United States) together with the High Sensitivity DNA Kit (Agilent Technologies, United States). Despite the adjustment of the index concentration, we detected a peak comprised of index dimers for some libraries and thus performed gel extraction on these libraries to separate the amplicon from the index dimers. Therefore, we ran the libraries for 3h at 65V on a 1.5% agarose gel, cut out the bands with the correct amplicon size and then used either the innuPREP Gel Extraction Kit (Analytik Jena, Germany) or the QIAquick Gel Extraction Kit (Qiagen, the Netherlands) according to the manufacturer’s protocol. A final quality control using the Agilent Bioanalyzer 2100 with the High Sensitivity DNA Kit was carried out before preparation of the libraries for high-throughput sequencing. Afterwards, all libraries were pooled into one tube equimolarly with initial amplicon concentrations ranging from 1.1 pmol/l to 9.4 pmol/l. The pooled sample was then run on an Illumina MiSeq® System (Illumina Inc, United States) with a MiSeq Reagent Kit v2 (500 cycles) according to the manufacturer’s protocol at the Max Planck Institute for Evolutionary Biology, Plön, Germany. The run on the Illumina MiSeq® System included 30 libraries stemming from our experiment and 6 belonging to another experiment. Index sequences that were used for multiple libraries could still be uniquely assigned to libraries, as the libraries differed in primer pairs used. To test for contamination in our workflow, we assigned negative controls to each plate and had at least one negative control per library. These controls confirmed that none of our libraries showed signs of contamination.

##### 2.3.2.2 Clean-up and processing of MHC

We de-multiplexed the raw data obtained from the illumina MiSeq runs using the AmpliSAS/AmpliCHECK genotyping tools developed by Sebastian et al. (2016). We first merged paired-end reads using AmpliMERGE and then removed low quality and erroneous sequences with AmpliCLEAN with a Phred quality score of 30. Demultiplexing and filtering with AmpliCLEAN resulted in 2,036,358 reads. We then used AmpliCHECK to explore the data and identify which advanced program parameters, such as minimum allele frequency and maximum number of alleles per amplicon, should be modified in the AmpliSAS step. Minimum allele frequency was estimated by comparing the frequency of artefacts with the frequency of putative alleles. We then tested different parameter settings and performed plausibility checks using preexisting pedigree data (Sanderson et al. 2015; Wells et al. 2018) that gave us information on the genotypes of parent-offspring triplets. For each exon we selected the median allele frequency of artefacts identified in AmpliCHECK as the minimum frequency threshold. We then determined the reproducibility among technical replicate samples using the formula [(Number of shared alleles*2)/sum of alleles in replicates] to identify the best settings for AMPLISAS (Table S3). After running AmpliSAS, we used AmpliCOMPARE to combine the results for technical replicates.

We then performed further manual quality and plausibility checks with the sequencing data obtained from the AmpliSAS pipeline described above. We first estimated the difference in bases between all sequences. We then used this data to follow the step-by-step procedure depicted in Figure S1. The number of base pairs that differed, the depth at which the sequence occurred, the number of samples the sequence appeared in, whether it appeared only together with a different, highly similar sequence, as well as whether it was present in both replicates of a sample were considered to identify a sequence variant either as a putative allele or an artefact. Sequences that were present in only one of the duplicate samples were kept for that individual if they fulfilled the criteria set for a putative allele in the step-by-step allele validation procedure described above.

Before manual allele validation, 70 sequences were retained for MHC-I exon 2, 16 sequences for MHC-I exon 3, and 23 sequences for MHC-II DRB exon 2. After clean-up following step-by-step procedure described in the previous paragraph, we were left with 37 sequences for MHC-I exon 2 (32 coding, 5 pseudogenes), 14 sequences for MHC-I exon 3 (all sequences are coding, but three of these have the same amino acid sequences as other sequences in the set, leaving 11 different amino acid sequences), and 17 for MHC-II DRB exon 2 (15 coding sequences, 2 pseudogenes).

We further used Megablast (Zhang et al. 2000) on NCBI GenBank to investigate similarity of the putative alleles to known carnivore sequences to validate the sequences. Next, we translated putative alleles, checked for stop codons and investigated whether conserved sites known to have structural importance can be found in the sequences (Kaufman et al. 1994; Reche and Reinherz 2003). We then used the results of the technical replicates to calculate reproducibility of the genotypes based on the formula [(Number of shared alleles*2)/sum of alleles in replicates].

We tried to assign alleles to loci using MHC typer V1.1 using a maximum-likelihood approach for reconstructing haplotypes while considering null alleles or copy number variation (CNV), identical alleles shared between loci as well as deviations from Hardy-Weinberg-Equilibrium (Huang et al. 2019), but were unsuccessful. Detailed methods for the approach used can be found in the supplementary material.

##### 2.3.2.3 Comparison of carnivore MHC diversity

Banded mongooses inbreed frequently, which may affect their levels of genetic diversity. To compare the total number of MHC sequences found in banded mongooses with the MHC population diversity of other carnivore species, we searched for studies reporting the total number of MHC sequences for MHC class I exon 2 and exon 3 as well as MHC class II DRB. We used the search engines Google Scholar and PubMed and the combined search terms “MHC”, “diversity” and “carnivore” (search performed: August 22^nd^ 2024). In addition, we searched for sequence similarity of our sequences in GenBank using Megablast (Zhang et al. 2000) and looked for linked literature for the carnivores that matched our MHC sequences. We identified 74 species and for each we compiled metadata on sample size, MHC class, number of exons amplified, and conservation status (Table S4+5). We assigned a conservation status (least concern (LC), near threatened (NT), vulnerable (VU), endangered (EN) or critically endangered (CR)) using the IUCN Red List of Threatened Species version 2024-1 (IUCN 2024) as a proxy for threat of extinction.

### 2.4 Statistical analyses

#### 2.4.1 Sequence polymorphism

We assessed sequence polymorphism through the number of polymorphic sites, average number of nucleotide differences, total number of mutations, nucleotide diversity, and pairwise identity using DnaSP 6.12.03 (Rozas et al. 2017). We estimated the amino acid p-distance with uniform rates in MEGA-X (Kumar et al. 2018). A detailed description of the methods can be found in Winternitz et al. (2023).

#### 2.4.2 Recombination

RDP4 v.4.101 software (Martin et al. 2015), which uses several distinct algorithms aimed at detecting recombinant sequences, was used to infer recombination signals in MHC-I exon 2 and 3 and MHC-II DRB exon 2. We used the following methods to infer recombination in our data set: RDP (Martin and Rybicki 2000), GENECONV (Padidam et al. 1999), BootScan (Salminen et al. 1995), SiScan (Gibbs et al. 2000), Maxchi (Smith 1992), Chimaera (Posada and Crandall 2001), and 3Seq (Boni et al. 2007). First, we screened the alignments of MHC-I and MHC-II nucleotide sequences for recombination with an automated exploratory search using the default settings, a statistical significance threshold of p = 0.05, and Bonferroni correction for multiple comparisons. We retained a recombination event for further analysis when it was detected by two or more algorithms. Second, we continued with a manual examination based on the guidelines of Martin et al. (2017). Therefore we sequentially examined all detected recombination events based on following criteria: the recombination event could be detected in more than one sequence, the characteristics of the recombination event (e.g. the identity of the recombinants and the breakpoint positions) could be verified, an average p-value across the recombinant sequences below 0.05 for the method detecting recombination, and no warning that the putative recombination signal has been caused by evolutionary processes other than recombination. A recombination event was accepted when all these criteria were fulfilled and rejected if this was not the case. The methods for recombination analysis have been described in Winternitz et al. (2023).

#### 2.4.3 Inference of selection

For both MHC classes, maximum likelihood fits for nucleotide substitution models were carried out in MEGA X (Kumar et al. 2018). We performed BIC-based model selection and found JC+G (G = 0.33, R = 0.5) to be the best fitting model for MHC-I exon 2. For MHC-I exon 3 JC+G also had the best fit (G = 0.10, R = 0.5) and for MHC-II DRB exon 2 we identified JC+I as the best model (G = 0.49, R = 0.5). Selection was determined based on comparing rates of nonsynonymous (dN) and synonymous (dS) substitutions. Positive (diversifying) selection is characterized by (dN>dS) driving changes in the amino acid sequence, whereas (dN<dS) represents negative (purifying) selection, and (dN=dS) implies neutral evolution.

We identified codon sites under natural selection by determining rates of dN and dS for each site based on two complimentary methods: FUBAR and MEME. FUBAR (Fast, Unconstrained Bayesian AppRoximation) infers rates based on a Bayesian approach (Murrell et al. 2013). It detects evidence of pervasive selection by assuming constant pressure.

Individual sites can experience different levels of positive and negative selection (episodic selection) and methods that can only detect pervasive selection will miss these effects. MEME (Mixed Effects Model of Evolution) uses a mixed-effects maximum likelihood approach instead to identify individual sites with signs of pervasive and episodic selection (Murrell et al. 2012). Positively selected sites (PSS) were thus those detected by either FUBAR (pervasive selection) or MEME (pervasive and episodic selection). Those detected by FUBAR only were classified as negatively selected sites. For analysis with FUBAR, posterior probability > 0.9 was set as our significance threshold, as these values strongly indicate natural selection. For analysis with MEME, our significance threshold was set to < 0.05. Both FUBAR and MEME selection inference was carried out on the Datamonkey server (Weaver et al. 2018, https://www.datamonkey.org/, last accessed: January 18, 2024).

We estimated mean dN-dS rates with standard error averaging over all sequence pairs using 1000 bootstrap replicates as a measure of strength of selection. For all classes and exons, analyses were carried out based on the Nei-Gojobori model with Jukes-Cantor correction for multiple substitutions (Nei and Gojobori 1986). All analyses determining the strength of selection as rates of (dN-dS) were performed in MEGA X (Kumar et al. 2018).To compare positions of the identified positively selected codon sites to those of the human PBR, the obtained amino acid sequences for MHC-I exon 2, exon 3, and MHC-II DRB exon 2 were aligned per exon and visually compared to PBRs identified in crystallographic analyses of human MHC molecules (Saper et al. 1991; Brown et al. 1993 please see Table S6). Since recombination can lead to overestimating positively selected sites, recombination inference was performed using RDP v.4.101 (Martin et al. 2015) before determining selection patterns. Inference of selection has been conducted according to the methods used in Winternitz et al. (2023).

#### 2.4.4 Phylogenetic relationships

To investigate the phylogenetic relationships among MHC sequences of banded mongoose and closely related carnivore species, we searched reference sequence genes (RefSeqGenes) available on NCBI GenBank (search performed on July 10, 2024). Sequences in this RefSeqGene set are intended to be well-supported, exist in nature, and represent a prevalent, ’normal’ allele. The sequence search used the key terms (“histocompatibility antigen”[All Fields] AND (“class I”[All Fields] OR “class II”[All Fields])) AND “carnivores”[porgn] AND “srcdb refseq”[Properties] AND alive[prop]). We selected MHC-I and MHC-II DRB genes from 12 carnivore species, five feliformia (cat-like) and seven caniformia (dog-like). These included the following families: Felidae (cats), Herpestidae (mongooses), and Hyaenidae (hyenas); Canidae (dogs), Ursidae (bears), Mustelidae (weasels, badgers, otters, and related species), Otariidae (sea lions and fur seals), and Phocidae (true seals); Feliforms: cheetah (*Acinonyx jubatus*), domestic cat (*Felis catus*), Canada lynx (*Lynx canadensis*), striped hyena (*Hyaena hyaena*), and meerkat (*Suricata suricatta*); Caniforms: dog (*Canis lupus*), giant panda (*Ailuropoda melanoleuca*), American black bear (*Ursus americanus*), Eurasian badger (*Meles meles*), domestic ferret (*Mustela putorius*), California sea lion (*Zalophus californianus*) and gray seal (*Halichoerus grypus*)). Human MHC-I genes and human and carnivore MHC-II DQB genes were selected to serve as outgroups to root trees. Since coding regions are more evolutionarily conserved than introns, we extracted the coding sequence (CDS) from each RefSeq gene for alignment and tree construction. We also included sequences of characterized MHC genes from the NCBI nucleotide database for Felis catus (Yuhki et al. 2008; Okano et al. 2020) and Acinonyx jubatus (Drake et al. 2004). Sequences of known nonclassical MHC genes (FLAI-A, FLAI-M, FLAI-O, FLAI-Q, FLAI-J, FLAI-L (Yuhki et al. 2008; Okano et al. 2020); DLA-12, DLA-64 (Burnett and Geraghty 1995); AIME-1906 (Zhu et al. 2013), HLA-E, HLA-F, HLA-G (D’Souza et al. 2019)) were included in the alignments to see if they clustered with banded mongoose sequences with strong support, suggesting homology and thus that we may have amplified nonclassical banded mongoose MHC sequences.

We aligned 107 sequences with 2109 nucleotide sites for MHC-I (Table S7) and 61 sequences with 831 nucleotide sites for MHC-II (Table S8) using MAFFT v7.490 (Katoh et al. 2002; Katoh and Standley 2013). Maximum likelihood phylogenetic trees for MHC sequences were created using the IQ-TREE webserver ((Trifinopoulos et al. 2016) http://iqtree.cibiv.univie.ac.at). We first used ModelFinder (Kalyaanamoorthy et al. 2017) to select the top supported model based on BIC (MHC-I best-fit model = TVM+F+R4; MHC-II best-fit model = GTR+F+R3), and then used IQ-TREE with 5000 ultrafast bootstrap (UFBoot) alignments to build the trees (Nguyen et al. 2015; Hoang et al. 2018). To compare the inferred MHC gene trees with the species tree, 100 trees based on the Mammal birth-death node-dated completed tree (Upham et al. 2019) were downloaded from https://vertlife.org/ for the 14 species in this study. Consensus topology and average branch lengths were computed with the consensus.edges function from the R package phytools v2.3-0 (Revell 2012) using 50% majority rule consensus. Phylogenetic tree figures were created using the R package ggtree v3.12.0 (Yu et al. 2017). Analyses and figures produced in R used version 4.4.1 (R Core Team 2023).

#### 2.4.5 Phylogenetic regression of carnivore MHC diversity

To compare the total number of MHC sequences found in banded mongooses with the MHC population diversity of other carnivore species, we conducted a Bayesian phylogenetic linear mixed model with the R package brms (Bürkner 2017), using a phylogenetic consensus tree estimated using the R package phytools (Revell 2024) of 100 phylogenetic trees (https://vertlife.org/phylosubsets/) dated by tip with mean branch lengths (Upham et al. 2019). Species not listed in the tree were assigned to the nearest relative to compute a phylogenetic covariance matrix including all species. To predict how total number of alleles (log_10_ transformed) was related to extinction risk, we included the fixed effects of conservation status (ordinal), log_10_ transformed sample size, the number of exons sequenced (1 or 2), MHC class (I or II), and the random effects of species to account for multiple observations per species and a covariance matrix of the relationship between species from branch lengths to account for non-independence among species. Our model used a gaussian distribution, default priors, 2 chains, 4000 iterations, 1000 warmup iterations, and the parameter adapt_delta increased to 0.99 to eliminate the number of divergent transitions during sampling. Model predictions were obtained using the R package ggeffects (Lüdecke 2018) and plotted using the R packages ggeffects, gginnards (Aphalo 2024) and ggplot2 (Wickham 2016).

## RESULTS

### 3.1 Genotyping results

All sequences that were verified as either classical or non-classical putative alleles can be found in the supplementary material.

#### 3.1.1 MHC-I exon 2

For MHC-I exon 2, 37 putative alleles of 228 bp length were detected (Fig. 1a). Apart from allele 17, all alleles had BLAST hits (E < 6e^-65^) with feliformia MHC-I exon 2 loci between 86.4% and 99.56% identity (please see supplementary files for detailed blast results). Allele 17 also had BLAST hits (E < 6e^-65^) but only with mammal species other than feliform species, with the highest sequence identity of 94.26% and E = 2^-90^ for the horse *Equus caballus*. Apart from five putative alleles (alleles 02, 04, 07, 22 and 30) all sequences translated without stop codons (on reading frame 1) and showed high conservation of sites known to be structurally important in other species’ classical MHC-I loci (Saper et al. 1991, see Table S9). To classify the putative alleles as classical or non-classical, we used the human MHC sequence (accession number AAA76608.2) as a reference, aligned it with our putative alleles and compared the nucleotides of the known conserved sites (Kaufman et al. 1994) with those from our putative alleles. Site Y84 from Kaufman et al. (1994) corresponded to site 108 in the HLA and banded mongoose alignment and was conserved in all putative alleles. In contrast site 83, which corresponded to HLA Y59, was not conserved in allele 05, 24, 26 and 37. These four alleles were thus classified as putatively nonclassical MHC sequences so were not considered for further selection analyses. The sequences used for the alignment can be found in Table S10.

**Figure 1.**
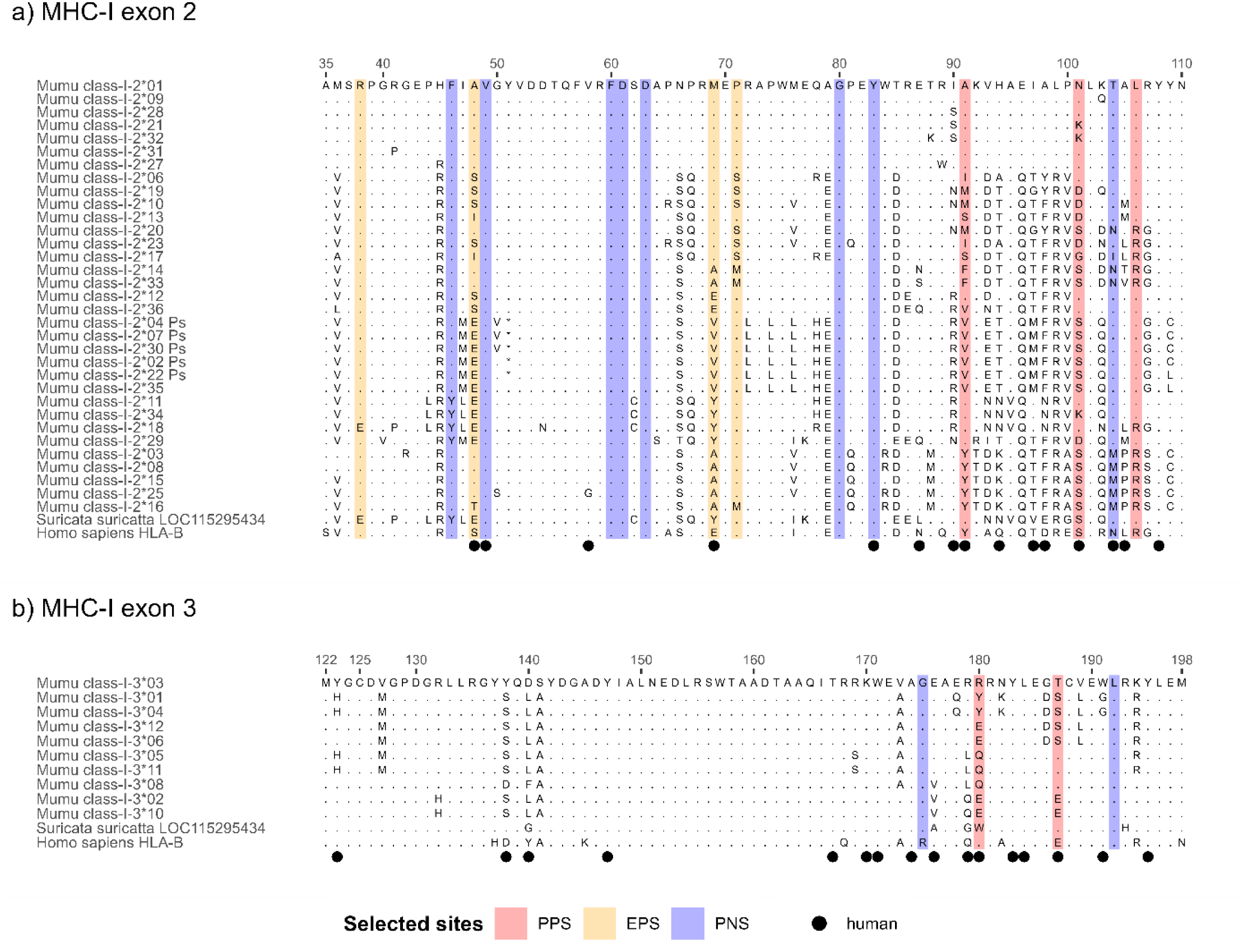
Alignment of amino acid sequences for classical MHC class I alleles of the banded mongoose (*Mungos mungo).* Displayed are the alignments for MHC-I exon 2 **(a)** aligned with HLA-B (accession number NC_000006, chromosome 6 assembly accession number NC_000006.12) and the meerkat (*Suricata suricatta*) MHC class I histocompatibility antigen sequence (accession number LOC115295434, genome NC_043706). Residues are numbered from 35 to 110 based on the HLA-alignment. MHC-I exon 3 amino acid sequences have been aligned with the same human and meerkat sequences as MHC-I exon 2, but the residue numbering established based on the HLA-sequence reaches from 122 to 198 **(b)**. Banded mongoose sequences are available on GenBank with the accession numbers PQ137681 - PQ137731. Dots represent sites that are identical to the amino acid shown at this site in the sequence on the top of the alignment. Circles indicate human peptide binding residues based on Saper et al. (1991). PPS: pervasive positive selection, EPS: episodic positive selection, PNS: pervasive negative selection. Pseudogenes were not included in the selection analysis but are shown for visualization purposes.

#### 3.1.2. MHC-I exon 3

For MHC-I exon 3, 14 putative alleles of 232 bp length were detected (Fig. 1b). All these alleles had BLAST hits (E < 1e^-5^) with feliform MHC-I exon 3 loci between 92.58% and 99.57% (please see supplementary files for detailed blast results). The alleles translated without stop codons (on reading frame 1) and showed high conservation of sites known to be structurally important in other species’ classic MHC-I loci (Saper et al. 1991, see supplementary files). Allele 01 and 04, allele 06 and 12, and allele 07 and 13 only contained synonymous substitutions and translated into the same amino acid sequence, respectively. Similar to MHC-I exon 2, we investigated whether putative alleles of exon 3 had conserved sites that were known from human MHC (Kaufman et al. 1994). HLA-A residue T143 from Kaufman et al. (1994) corresponded to site 167 of the HLA/banded mongoose alignment and was conserved in all putative alleles (Table S9). In contrast Y171, which corresponded to site 195 in the mongoose sequence, was not conserved in allele 07, 09, 13 and 14. These four alleles were thus classified as non-classical MHC sequences so were not considered for further selection analyses. The sequences used for the alignment are listed in Table S11.

#### 3.1.3. MHC-II DRB exon 2

For MHC-II DRB exon 02, 17 putative alleles of 201 bp length were detected (Fig. 2). Apart from allele 05 and 16 all alleles had BLAST hits (E < 1e^-5^) with feliform MHC-II loci between 86.53% and 95.52% identity (please see supplementary files for detailed blast results). Allele 05 was most similar to the sequence of the bamboo lemur *Hapalemur griseus griseus* (E = 1^-71^, % ident. = 92.86) and allele 16 had the best match with a sequence from the Himalayan bear Ursus thibetanus (E = 2^-64^, %ident. = 91.1). Except for allele 08 and 13, all alleles translated without stop codons (on reading frame 3). The sequences showed high conservation of sites known to be structurally important in MHC-II sequences of other species’ classic MHC-II loci (Brown et al. 1993; Singh 1998, see supplementary files). We investigated whether putative alleles of DRB exon 2 had conserved sites that were known from human MHC (Kaufman et al. 1994). Conserved site H81 from Kaufman et al. (1994) corresponded to site H110 in the HLA sequence retrieved from GenBank (accession number NP002115.2) and the aligned mongoose sequence and was conserved in all putative alleles (Table S9). In contrast W61, which corresponded to site 90 in the HLA/mongoose sequence alignment, was not conserved in allele 07 and 11. These two alleles were thus classified as non-classical MHC sequences that were not considered for further selection analyses. The sequences used for the alignment are depicted in Table S12.

**Figure 2.**
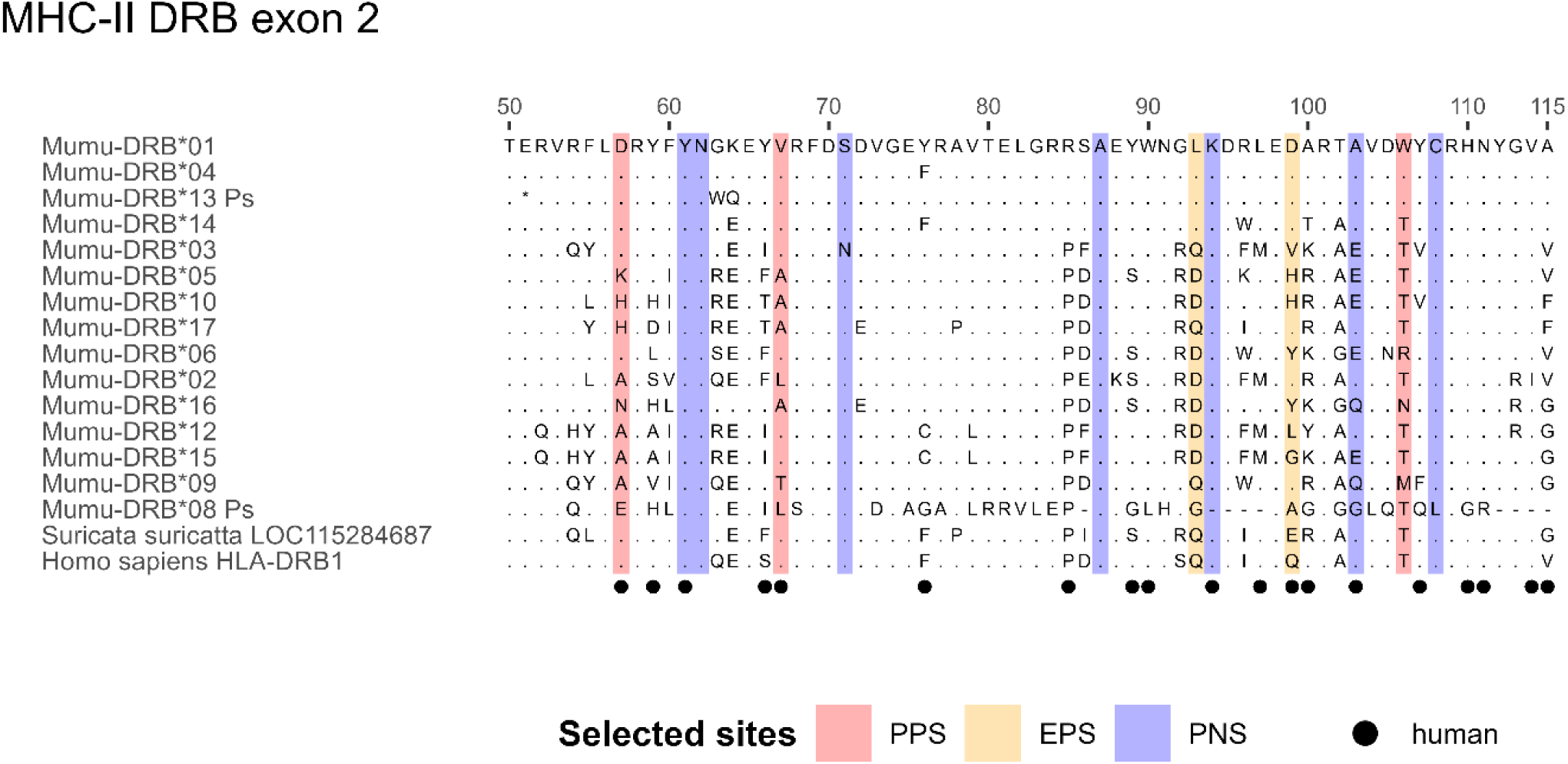
Alignment of amino acid sequences for classical MHC class II DRB exon 2 alleles of the banded mongoose (*Mungos mungo).* Alignments were made with HLA-DRB1 (genome accession number NC_000006) and DLA class II histocompatibility antigen DR-1 beta chain-like of the meerkat (*Suricata suricatta*) (accession number LOC115284687, genome accession number NC_043706). Residues are numbered from 50 to 115 based on the HLA-alignment. Banded mongoose sequences are available on GenBank with the accession numbers PQ137732 - PQ137748. Dots represent sites that are identical to the amino acid shown at this site in the sequence on the top of the alignment. Circles indicate human peptide binding residues based on Brown et al. (1993). PPS: pervasive positive selection, EPS: episodic positive selection, PNS: pervasive negative selection. Pseudogenes were not included in the selection analysis but are shown for visualization purposes.

### 3.2 Assigning alleles to loci

We tried to assign alleles to loci for MHC class I and II but were unable to establish the same assignment with the same optimal BIC over multiple runs. We also acknowledge that henceforth we use the term ‘allele’ to designate sequence variants from unknown loci. MHC nomenclature from this study takes the first 2 characters of the genus species name (Mumu) following the recommendations of Maccari et al. (2018). For class I, the four-letter species abbreviation is followed by class-I and the respective exon, either -2 or -3. The last part of the sequence nomenclature for both classes is the sequence number, ordered by decreasing population frequency. Finally, the abbreviation Ps (pseudogene) indicates non-coding sequences.

### 3.3 Individual allelic diversity

For MHCI-I exon 2, 321 samples were successfully genotyped with a mean reproducibility of 89.4%. Read number per sample ranged from 100 to 5000 reads, with a mean of 637 reads per sample. Allele numbers per individual ranged from 3 to 14 alleles with a mean of 7.56. Three individuals carried more than 12 alleles: BF773, BM842, and BM212, but all alleles passed filtering based on our allele validation criteria. Individual BF773 carried 13 alleles of which seven were found in both technical replicates. Of the remaining 6 alleles, two could be confirmed in one of the parents for which a genotype was available. The four alleles that were only found in one replicate were frequent alleles (found in 122, 122, 93 and 38 individuals) and can thus be considered true alleles. Individual BM842 carried 13 alleles of which 10 were found in both technical replicates. The remaining three alleles that were not present in both technical replicates of BM842 were present in 93, 37, and 24 individuals in total. Thus, none of the alleles were rejected. Individual BM212 carried 14 alleles of which five were found in both technical replicates. Of the nine alleles found only in one replicate, six alleles were found in three siblings and a further allele was found in the father. The two remaining alleles were frequent alleles (found in 87 and 55 individuals) and were therefore considered true alleles.

For MHC-I exon 3, 285 individuals were successfully genotyped with a mean reproducibility of 96.6%. Read number per sample ranged from 110 to 5000 reads, with a mean of 1215 reads. Allele numbers per individual ranged from 1 to 5 with a mean of 1.6. Two individuals carried more than 4 alleles, but the alleles could all be confirmed based on our validation criteria. Individual DF061 carried five alleles with all alleles appearing in both technical replicates. Individual HM313 carried five alleles, with three of these alleles appearing in both technical replicates. A further allele was found in three siblings, and the final was the most frequent allele (found in 280 individuals) so was not rejected.

For MHC-II DRB exon 2, 384 individuals were successfully genotyped with a mean reproducibility of 97.7%. Read numbers varied from 101 to 5000 reads per sample with a mean of 1224 reads. Individuals had between 1 and 6 alleles, with a mean of 2.49. There was one individual, 8K, with 6 alleles, 5 of which were found in the technical replicate. The remaining allele was found in an offspring of 8K and thus confirmed as well.

The frequencies of the putative alleles strongly differ between the genes investigated (Fig. 3). Whereas MHC-II DRB exon 2 genotypes are dominated by coding sequences (Fig. 3c), MHC-I exon 2 genotypes include non-coding sequences at a much higher rate (Fig. 3a). The two pseudogenes Mumu class-I-2*02-Ps and -04-Ps appear in more than half of the individuals in this study. In contrast, Mumu-DRB*08-Ps appears in a tenth of the tested individuals and Mumu-DRB*13-Ps only in 1% of the samples. For MHC-I exon 3 no non-coding sequences were identified (Fig. 3b).

**Figure 3.**
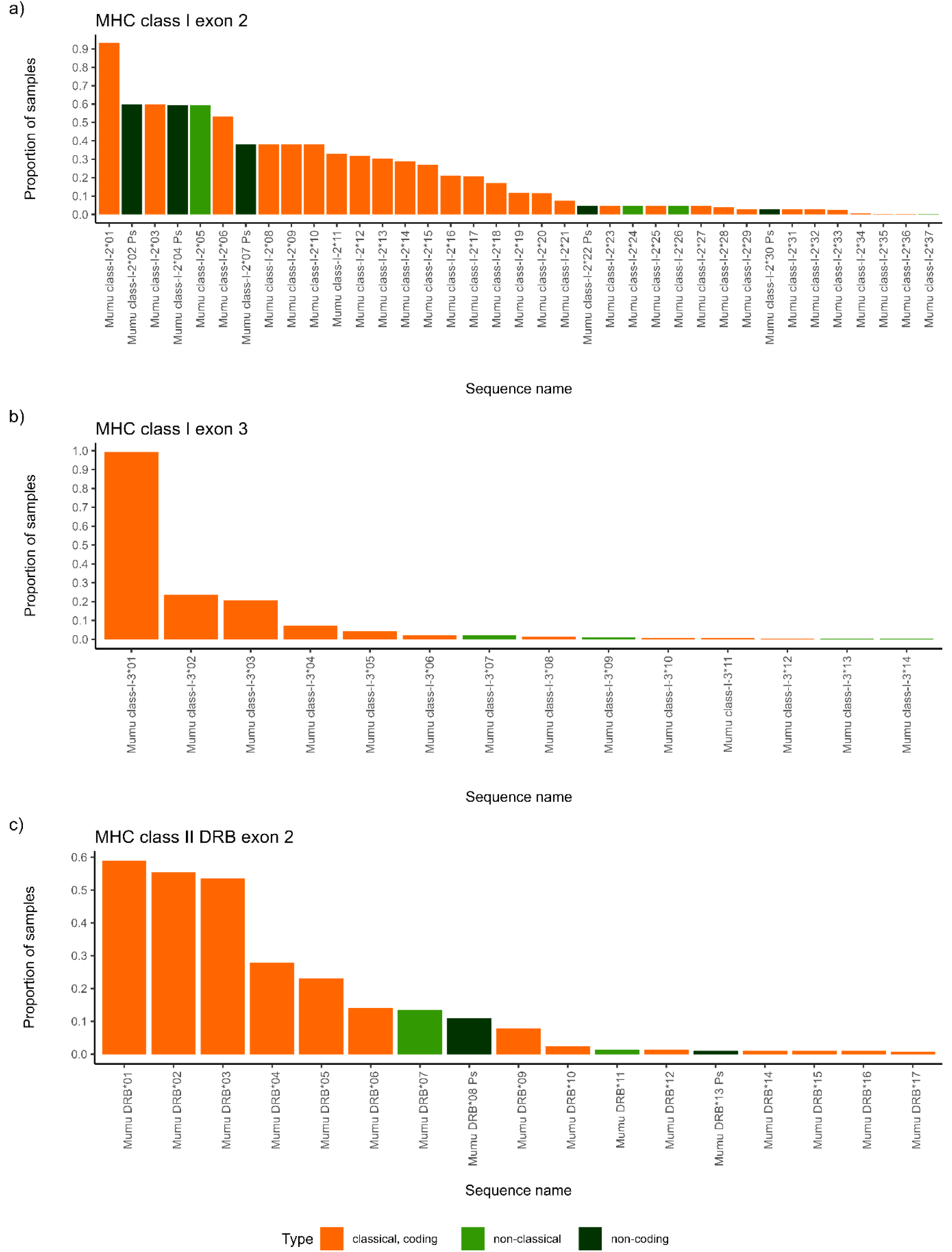
Allele frequencies for MHC-I and -II. The frequency with which the identified putative alleles appear in the samples individuals is displayed. Alleles are either identified as non-classical (dark green) or classical sequences (orange) for MHC-I exon 2 **(a)**, for MHC-I exon 3 **(b)** and for MHC-II DRB exon 2 **(c)**. Within the classical alleles, non-coding sequences are highlighted (light green).

### 3.4 Sequence polymorphism

MHC-I exon 3 had the highest number of nucleotides with 232, followed by MHC-I exon 2 with 228 nucleotides and MHC-II DRB exon 2 with 201 nucleotides. For MHC-I exon 2, 28 different alleles were found with 100 polymorphic sites and 132 mutations in total (Table 1). The average number of nucleotide differences was k = 32.238 and the nucleotide diversity Pi = 0.1414. The mean amino acid p-distance was 0.251 ± 0.029. For MHC-I exon 3, diversity was overall lower with 10 distinct alleles, 30 polymorphic sites and 36 mutations detected. The nucleotide diversity for MHC-I exon 3 was Pi = 0.0565 and the mean amino acid p-distance equaled 0.111 ± 0.025. For MHC-II DRB exon 2, 13 distinct alleles were detected with 79 polymorphic sites and 105 mutations. The nucleotide diversity was Pi = 0.1544 and the mean amino acid p-distance was 0.259 ± 0.037. Pairwise identity of MHC-I exon 2 was 85.3% and for exon 3 94.2%, which corresponds to a divergence of 14.8% and 5.8% respectively. For MHC-II DRB exon 2 pairwise identity was 84.3% resulting in a divergence of 15.7%. Consequently, both MHC classes and all exons fulfill the minimum requirement of 5% sequence divergence and thus the power required for the detection of recombination by most methods (Posada and Crandall 2001).

**Table 1.** Sequence polymorphism of banded mongoose MHC-I and MHC-II. S: number of polymorphic sites, Eta: total number of mutations, k: average number of nucleotide differences, Pi: nucleotide diversity, AA p-distance: number of amino acid differences per site.

| MHC class | Exon | Nucleotide number | Allele number | S | Eta | K | Pi | AA p-distance |
| --- | --- | --- | --- | --- | --- | --- | --- | --- |
| MHC-I | 2 | 228 | 28 | 100 | 132 | 32.238 | 0.1414 | $0.251 \pm 0.029$ |
| MHC-I | 3 | 232 | 10 | 30 | 36 | 13.111 | 0.0565 | $0.111 \pm 0.025$ |
| MHC-II | DRB 2 | 201 | 13 | 79 | 105 | 31.026 | 0.1544 | $0.259 \pm 0.037$ |

### 3.5 Recombination

For both MHC classes and all exons no recombination event met our validation threshold. This means that according to our criteria no sequence is considered a recombination product of other sequences since no breakpoints were confirmed by the algorithms used by GARD or RDP4.

### 3.6 Inference of selection

MHC-I exon 2 showed stronger signs of selection compared to MHC-I exon 3 and MHC-II DRB exon 2 (Tab. 2). FUBAR and MEME detect pervasive and episodic selection respectively and found 7 sites under positive selection for MHC-I exon 2 compared to 5 in MHC-II DRB exon 2 and 2 in MHC-I exon 3. A similar pattern can be observed for negative selection, which was detected for 8 sites in MHC-I exon 2, for 7 in MHC-II DRB exon 2 and for 2 sites in MHC-I exon 3. Negatively selected sites mostly showed no polymorphism at all and were fixated for a single amino acid, e.g. at site 60, 61 and 63 for MHC-I exon 2, at site 192 for MHC-I exon 3 and at site 61 and 62 for MHC-II DRB exon 2. However, there were also polymorphic sites under negative selection, e.g. site 103 of MHC-II DRB exon 2 that we also identified to represent a PBR site in the HLA-sequence. Interestingly MHC-I exon 3 was fixed for a G (glycine, hydrophobic) in banded mongooses as well as meerkats but the HLA-B sequence showed an R (arginine, basic) at this site.

**Table 2.** Patterns of selection detected in MHC class I exon 2 and 3 and MHC class II DRB exon 2 of banded mongooses. dN-dS: ratio between non-synonymous and synonymous describing the direction of selection, PSS: positively selected site, PBR: peptide binding region.

| MHC class | Exon | n<sub>alleles</sub><br>(n<sub>AAsites</sub>) | Number of sites under selection |  | All sites |  | PSS |  | Human PBR |  |
| --- | --- | --- | --- | --- | --- | --- | --- | --- | --- | --- |
|  |  |  | Positive | Negative | dN - dS | P | dN - dS | P | dN - dS | P |
| MHC-I | 2 | 28 (76) | 7 | 8 | 0.908 | 0.183 | 5.700 | 4*10<sup>-8</sup> | 0.917 | 0.180 |
| MHC-I | 3 | 10 (77) | 2 | 2 | 1.129 | 0.131 | 2.402 | 0.009 | 0.164 | 0.051 |
| MHC-II | DRB 2 | 13 (66) | 5 | 7 | -0.042 | 1 | 5.146 | 1.6*10<sup>-7</sup> | 0.281 | 0.390 |

The strength of selection defined as the ratio of non-synonymous vs. synonymous mutations didn’t show a clear pattern (Tab. 2), with no significant selection found when considering all sites, nor for sites corresponding to the human PBR. However, when considering the positively selected sites (PSS) only, significant positive selection could be detected for both classes and all exons.

### 3.6 Phylogenetic analysis

To investigate the phylogenetic relationships among MHC sequences of banded mongooses and related carnivore species, we created MHC gene sequence trees and compared these with the species tree. We found that sequences mostly clustered with high (≥ 95%) probability within species and within families (Felidae (cats), Herpestidae (mongooses), and Hyaenidae (hyenas); Canidae (dogs), Ursidae (bears), Mustelidae (weasels, badgers, otters, and related species), Otariidae (sea lions and fur seals), and Phocidae (true seals)) (Fig. 4). The suborders feliformia (cat-like species) tended to cluster independently, with exceptions for MHC-I including a clade with a known nonclassical cat locus (FLAI-A) and predicted nonclassical loci in the Eurasian badger and domestic ferret (Patr-E-like). The exceptions to clustering by suborder feliformia for MHC-II were the DQB and DRB locus clades. Therefore, within our trees there are some homologous MHC-I loci shared among carnivores but most evolved after divergence at the family level. MHC-II loci appear to have longer evolutionary history than MHC-I, with the DQB clade including sequences from a primate and carnivores, the DRB clade spanning carnivores, and orthologous loci clustering by family.

**Figure 4.**
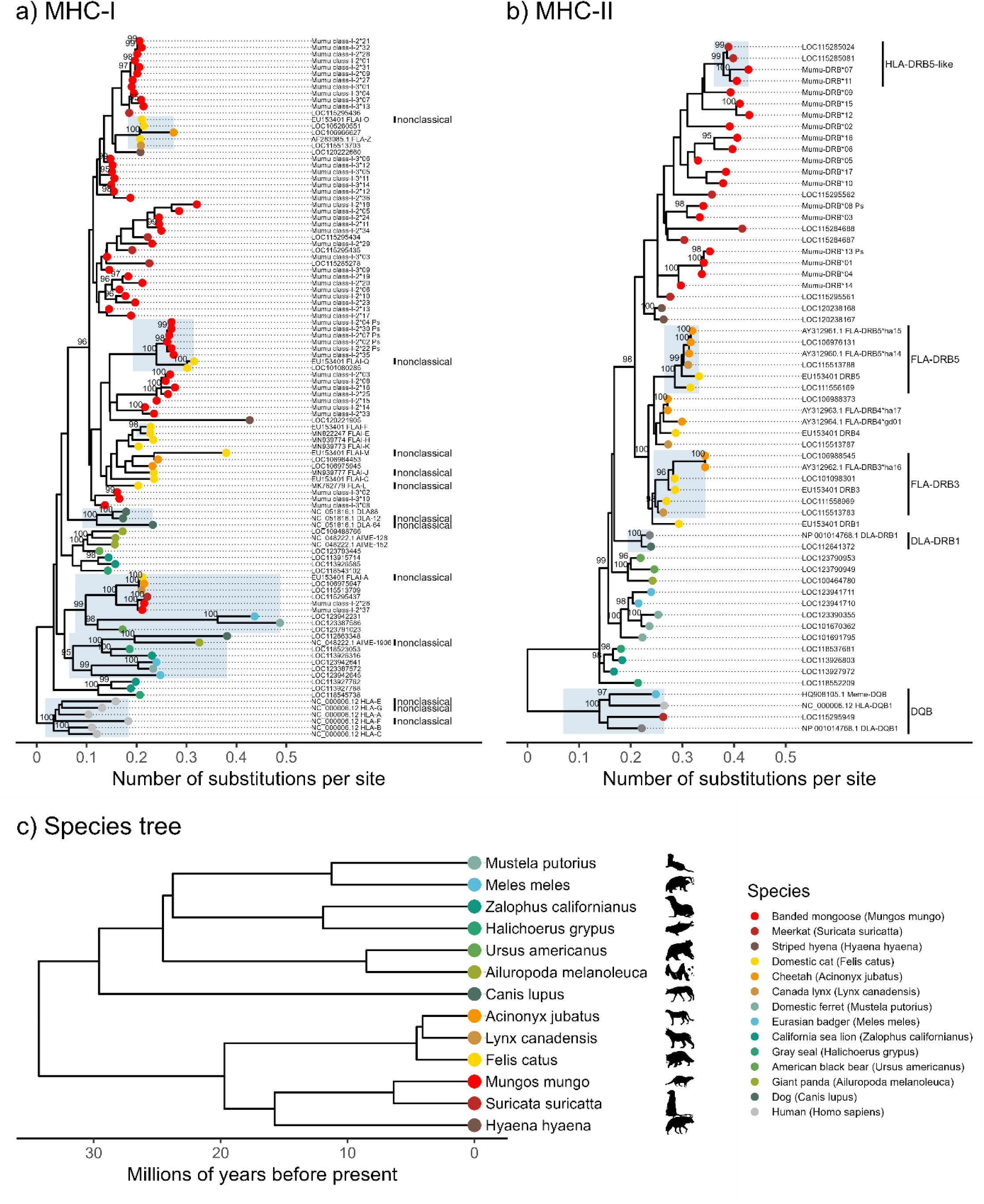
Phylogenetic relationships of banded mongoose MHC-I and MHC-II sequences across carnivores. Maximum-likelihood phylogenetic trees were created for banded mongoose MHC sequences using RefSeq CDS of MHC alleles from diverse carnivore species. UFBoot support values greater than 95% are shown―the recommended cutoff for high probability that a clade is true. **(a)** A phylogenetic tree including 37 and 14 *Mungos mungo* sequences of MHC class I exon 2 and 3, respectively. The tree was rooted using the human HLA class I allele clade as an outgroup. Clades with strong support that include known nonclassical loci are highlighted. Banded mongoose sequences cluster both interspecifically and by species or with the closest relative, the meerkat. **(b)** A phylogenetic tree including 17 *Mungos mungo* sequences of MHC class II DRB exon 2, rooted by a clade of MHC-II DQB sequences. Known locus-specific clades with strong support are highlighted, illustrating that MHC-II loci tend to have deeper evolutionary history than MHC-I loci, with duplicated MHC-II loci shared across species. **(c)** The phylogenetic consensus tree of 13 species included in the MHC sequence trees. Tips are color-coded so that feliformia (cat-like carnivores) have warm colors (red-yellow) and caniformia (dog-like carnivores) have cool colors (blue-green). Artwork from phylopic.org.

Banded mongoose MHC-I sequences mostly clustered by species or with meerkats, but one well-supported clade included noncoding sequences and the cat nonclassical locus FLAI-Q, and another included putative nonclassical sequences Mumu class I-N*26 and 37 and the cat nonclassical locus FLAI-A (Fig. 4a). Clades with strong support that include known nonclassical loci (FLAI-O, FLAI-Q, FLAI-A, AIME-1906) are highlighted to show that these include multiple species and could suggest homology or convergent evolution. Other known nonclassical loci (FLAI-F, DLA-12, DLA-64, HLA-E, HLA-F, HLA-G) cluster with classical loci within species, suggesting species-specific gene duplication and subsequent divergence. Thus, banded mongooses may have both homologous (interspecific) and orthologous (species-specific) nonclassical loci, as predicted by the conserved residue motifs described in section 3.1.1-3.1.3.

For MHC-II, banded mongoose sequences form a monophyletic clade with meerkats and striped hyena, suggesting MHC DRB loci from these species were present in a common ancestor approximately 17.7 million years ago (Upham et al. 2019). Meerkats appear to have six DRB loci based on incomplete genomic scaffolds (NCBI RefSeq assembly: GCF_006229205.1). Banded mongooses have at least four DRB loci based on a maximum of seven NGS sequences recorded per individual, though we were not able to use phylogenetic inference to identify individual loci from our 202-205 bp sequences.

### 3.7 Carnivore MHC diversity

To investigate whether our sample size was sufficiently large to capture the majority of the allelic diversity of our study population, we conducted an accumulation analysis using the package vegan version 2.6-8. The accumulation curves (Fig. S2a-c) showed that the samples sizes used in this study were sufficient to capture the vast majority of allelic diversity. We thus do not think that the sample size led to underestimation of allelic diversity.

To compare banded mongoose population MHC allelic diversity with those of other carnivore species, we needed to control for sampling effort, phylogenetic relationships, and population size. We compiled 74 observations of 39 carnivore species with MHC sequence data (consensus phylogenetic tree as Fig. S3). The phylogenetic signal estimate was very low (lambda=0.02 95% CI [0-0.10]), indicating there was little effect on MHC allelic diversity attributed to phylogenetic relatedness. Controlling for species and phylogenetic relatedness, we found as expected that the number of MHC alleles genotyped increased with sampling effort (posterior probabilities =0.25, 95% CI[0.14 - 0.36]), that there were fewer alleles genotyped for MHC-II than MHC-I (-0.35, [-0.50 - -0.21]), and that the number of alleles showed a significant linear decreasing trend as threat of extinction increased (Fig. 5). We also found no evidence that banded mongooses have lower MHC diversity at MHC-I or MHC-II when accounting for sampling effort and conservation status. Banded mongoose total allele numbers per conservation status “least concern” were well within the 5^th^ and 95^th^ percentiles (MHC-I, 28 alleles = 75%; MHC-II, 13 alleles = 54%). In contrast, the 1% and 99% outliers belonged to the Siberian tiger with 1 MHC-II allele from 1 individual and the raccoon with 66 MHC-II alleles from 246 individuals.

**Figure 5.**
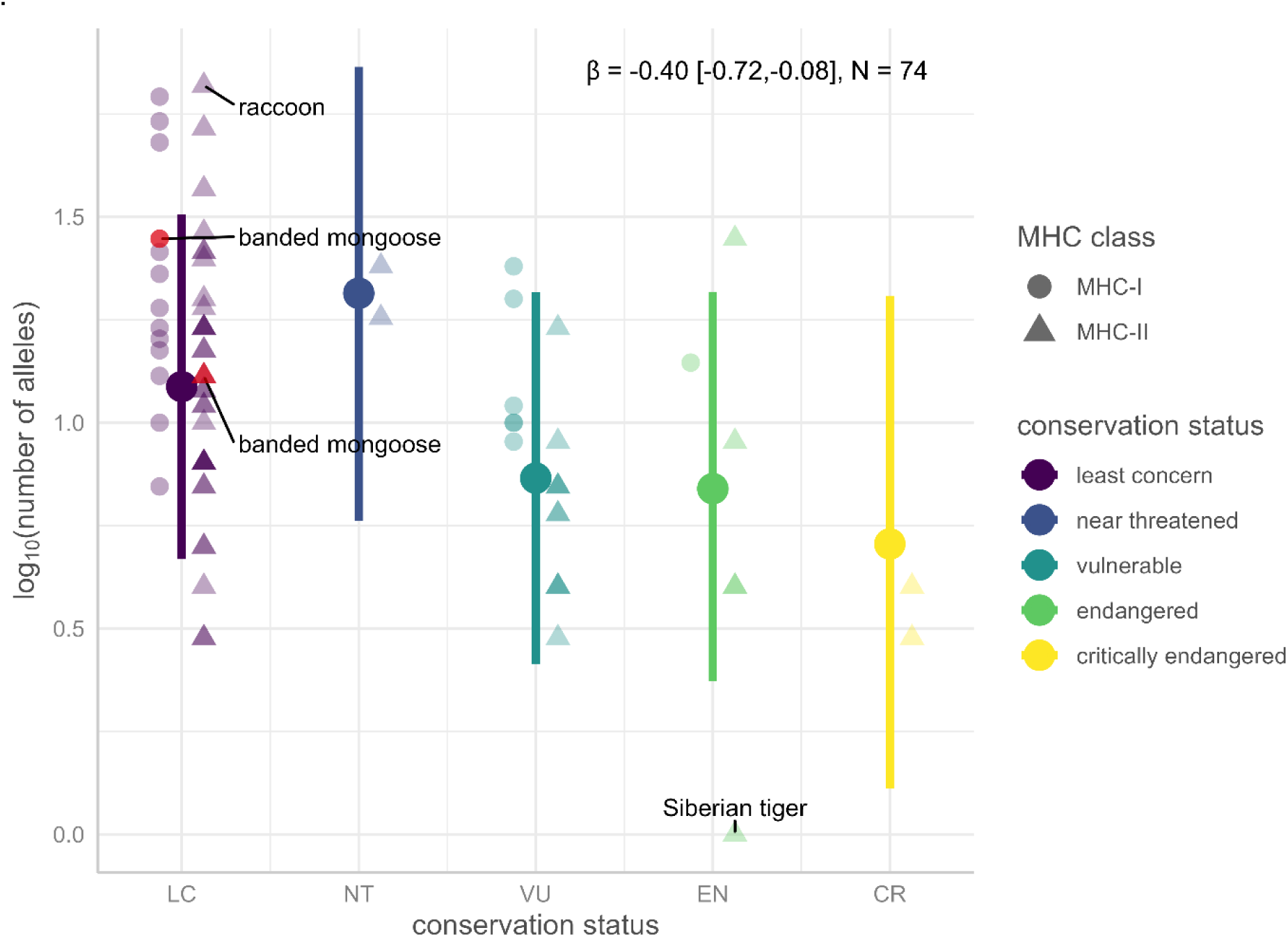
Predicted number of MHC alleles decreases with threat to conservation status. Predictions shown as colored points with 95% confidence intervals come from a Bayesian phylogenetic linear mixed model, controlling for species, phylogenetic relatedness, MHC class and sampling effort (numbers sampled and exons sampled). Raw data are shown for MHC-I and MHC-II separately. Banded mongoose values are within the 95% CI range for species of least concern, highlighted in red. Values below the 1% and above the 99% range are labeled by species.

## DISCUSSION

The banded mongoose shows MHC polymorphism and selection patterns similar to those seen in other species. Remarkably, despite their high inbreeding levels, their MHC diversity is comparable to that of other species not at risk of conservation concern.

Comparison of MHC population diversity within and between different mammalian species has to be done with caution, as concluded by Kelley et al. (2005) and emphasized by Castro-Prieto et al. (2011). Demographic history of a population, the degree of admixture, as well as selective pressures can affect MHC diversity. Further factors influencing the diversity and polymorphism detected are the methods used to retrieve the putative alleles, the number of genes investigated, the number of populations sampled as well as the total sample size (Castro-Prieto, Wachter, and Sommer 2011). For that reason, we performed phylogenetic mixed model regression accounting for the conservation status, the sample size, the number of exons sequenced, the MHC class and the species itself. The general pattern showed that MHC diversity decreased with increasing conservations threat status. This indicates that species suffering from severe population decline have lower MHC diversity compared to less threatened species, which can be detrimental to their conservation (Sommer 2005; Ujvari and Belov 2011; Coker et al. 2023). When looking at banded mongooses, a species of least conservation concern but with low levels of dispersal and frequent inbreeding, MHC diversity is not lower than expected for a wild carnivore of least conservation concern. This is surprising given that two thirds of the population result from inbreeding and 7.1% stem from full-sibling or parent offspring matings (Wells et al. 2018).

This could indicate the effective population size is large and less impacted by genetic drift that would decrease MHC genetic diversity. Alternatively, MHC diversity may be preserved by balancing selection acting through mate choice. MHC-dependent mate choice has been investigated in a wide range of species and mate choice for MHC-diverse mates is common (Kamiya et al. 2014; Winternitz et al. 2017; Winternitz and Abbate 2022). Another possibility could be that parasite-mediated balancing selection preserves MHC diversity. Since the MHC is a key player in the adaptive immune response, higher MHC diversity could result in greater parasite resistance (Hughes and Nei 1989; Takahata and Nei 1990; Pierini and Lenz 2018). To effectively determine whether MHC diversity is maintained through balancing selection despite genetic drift, comparison between neutral genetic diversity and MHC diversity is necessary. One way to achieve this is by investigating links between MHC diversity and neutral markers, such as microsatellites or by comparing MHC heterozygosity with overall genomic heterozygosity. Future studies should thus investigate correlations between neutral and MHC diversity as well as parasite load and fitness proxies such as survival and reproductive success in relation to MHC diversity, and analyze patterns of mate choice.

The number of putative alleles found for the two classes in banded mongooses differed, with MHC-I exon 2 having the highest number of sequences retrieved, followed by MHC-II DRB exon 2 and finally MHC-I exon 3 showing the lowest number of sequences. This discrepancy in the total allele number between the two exons of class I might be caused by the different selection pressures acting on them. In line with this, we found higher levels of positive selection in class I PSS for exon 2 compared to exon 3, which has been observed in other species such as the cheetah (*Acinonyx jubatus*) (Castro-Prieto, Wachter, and Sommer 2011) and the Eurasian badgers (genus: *Meles*) (Abduriyim, Nishita, et al. 2019). This difference in selection acting on the two exons is likely caused by the structure of the MHC-I binding pocket and the involvement of the two exons in forming it (Zhang et al. 1998). Exon 2 and 3 each build one half of the peptide binding groove (Saper et al. 1991) of the class I molecule and, because of the different structural involvement in the forming the peptide binding groove, positive selection may act more strongly on exon 2 compared to exon 3 (Zhang et al. 1998). The PBR of the MHC-I molecule is made up of the α1- and α2-domain, which are encoded by exon 2 and 3 respectively and form pockets that bind the peptide side chains. Whereas under the α1-helix there are three pockets situated that bind the peptide’s side chains, along the α2-helix there lies only one pocket (Zhang et al. 1998). These multiple binding pockets for the α1-domain might lead to selection acting more strongly on exon 2 compared to exon 3, as also observed in cheetahs and suggested by Castro-Prieto et al. (2011). These molecular mechanisms causing certain selection signatures could have led to exon 2 being more polymorphic in the population compared to exon 3. Alternatively, the primers we used may have had lower amplification success for MHC-I exon 3 than for exon 2.

Selection patterns differed for the different classes and exons, too. MHC-I exon 2 showed the highest dN-dS ratios for the PSS followed by MHC-II DRB exon 2. The cause for this might be the exons’ different role in pathogen binding (Klein 1986; Neefjes et al. 2011; Rammensee et al. 2013), that causes selection to be stronger and maintain diversity differently in different loci. The pattern observed in humans shows a higher signature of positive selection in MHC-I compared to MHC-II loci (Satta et al. 1994). The large number (5 for MHC-I exon 2) of noncoding sequences (pseudogenes) might further indicate that MHC-I exon 2 is under birth-and-death evolution (Takahashi et al. 2000; Kuduk et al. 2012; Abduriyim, Zou, et al. 2019). During birth-and-death evolution gene duplication creates new genes that can either persist for a long time or lose function because of deleterious mutation and end up as pseudogenes (Nei et al. 1997). However, since we looked at the two halves of the peptide binding groove of class I separately by sequencing both exon 2 and 3, this comparison must be done with caution, as the signatures of selection differed greatly for the two exons and no solid conclusion can be drawn without linking the diversity of the two exons of MHC-I for a comparison with MHC-II. To allow a solid comparison, future studies should investigate the diversity of the complete PBR for both classes combined, as suggested by Winternitz et al. (2023), since comparing PBR halves separately can bias the outcome substantially.

As expected, the sites identified as sites of positive selection showed higher levels of dN-dS ratios compared to the signature of selection in the full sequences for both classes and all exons. Studies in other mammals, such as the Eurasian beaver (*Castor fiber*) (Babik et al. 2005), and carnivores such as Namibian leopards (*Panthera pardus pardus*) (Castro-Prieto, Wachter, Melzheimer, et al. 2011), cheetahs (*Acinonyx jubatus*) (Castro-Prieto, Wachter, and Sommer 2011), Bengal tigers (*Panthera tigris tigris*) (Pokorny et al. 2010), found higher rates of non-synonymous mutations compared to rates of synonymous mutations for the PBR compared to the whole sequence. Interestingly, assuming that the human PBR sites also represent the PBR in banded mongooses, the dN-dS rates at those sites did not reproduce the patterns of positive selection being stronger compared to the complete sequence. This missing pattern for the human PBR sites could be attributed to these sites not being as evolutionary conserved as other functionally important sites that could be identified for the classical sequences found in this study. It is rather unlikely to be the main cause though, since previous studies have found the PBR of the human to be structurally similar in felids (Yuhki et al. 1989; Yuhki and O’Brien 1994).

The MHC gene sequence trees with related carnivore species showed that the suborder feliformia mostly clustered independently from caniformia. MHC-I loci sequences mostly clustered within species or families and MHC-II DRB and DQB loci clustered among more distant taxa, suggesting that duplication events at MHC-I loci happened more recently than at MHC-II loci and may be species specific. These patterns observed indicate that MHC-II loci have longer evolutionary history than MHC-I, which is in line with previous findings on the evolutionary trajectory of the two classes (Okamura et al. 2021).

Phylogenetic inference suggests that nonclassical loci are shared among felids (cats, mongooses, meerkats) thereby indicating trans-species polymorphism. But nonclassical loci are also found within species, e.g. FLAI-J, L. Molecular evidence of the comparison of conserved sites of HLA-sequences with the sequences recovered in this study suggests that banded mongoose may have both nonclassical MHC-I and II loci. These findings need to be considered before testing for associations between allelic diversity and fitness because nonclassical loci are typically not involved in antigen binding (Braud et al. 1999; Alfonso and Karlsson 2000; Adams and Luoma 2013).

Both the phylogenetic trees and the number of putative alleles found per individual indicate that banded mongooses have at least 4 MHC-II loci. However, we were not able to infer the number of loci from our data with the current methods despite relying on a thorough validation pipeline, a well-established NGS protocol with technical replicates of every sample, negative controls on each plate, and plausibility checks with pedigree data and residues with structural importance. This could be caused by large differences in the number of sequences recovered per individual. Causes for such large variation could be either copy-number variation (like in cats *Felis catus* (Yuhki et al. 2008)) or amplification bias during PCR. Again, assignment of the putative alleles to loci could help reveal patterns of copy-number variation as well as enable estimating heterozygosity levels, determining whether alleles are shared across loci, and investigating potential inter-locus gene conversion. Amplification bias was anticipated when designing this study, and we tried to avoid uneven distribution of reads by using a normalization kit. However, this only prevents amplification bias between samples but not between loci. Locus-specific primers developed from fully resolved banded mongoose genomes would answer this question, and this is a goal for the near-future.

## CONCLUSION

The high throughput sequencing data revealed that banded mongooses show distinct patterns of polymorphism and selection for the two halves of the binding groove of MHC-I built by exon 2 and 3, thus our study provides valuable information about the selection pressure acting on the different parts of the peptide. Phylogenetic analysis and comparison of the putative alleles with known conserved and structurally important sites of classical MHC loci indicated that banded mongooses have nonclassical loci for both MHC classes. These nonclassical class I sequences clustered with nonclassical sequences of other carnivore species, thereby indicating convergent evolution and trans-species polymorphism. Furthermore, nonclassical sequences clustered with classical ones, suggesting that species-specific gene duplication caused this pattern. Moreover, a general pattern of decreasing MHC diversity with increasing conservation threat status was observed. Most strikingly, the comparison of MHC diversity with other carnivores of least conservation concern showed that, despite high levels of inbreeding within the population, banded mongooses have comparable MHC diversity. This might be caused by parasite-mediated balancing selection or MHC-based mate choice causing MHC-diversity levels to persist despite frequent inbreeding. Future studies should aim at linking these patterns of historical selection with evidence for current selection through mate choice, reproductive success, and survival.

## Supporting information

all sequences_MHCI-exon2

all sequences_MHCI-exon3

all sequences_MHCII-DRB-exon2

Blast results - MHCI exon 2

Blast results - MHCI exon 3

Blast results - MHCII DRB exon 2

## Acknowledgements

We would like to thank the field crew of the banded mongoose research project and all the other PhD students and Postdocs that have been involved in collecting the valuable data and tissue samples. We also would like to thank Sven Künzel for Illumina Sequencing and Tobias Krause and Yvonne Kutter for sharing their expertise and letting us use their lab equipment.

## Authors’ contributions

NS contributed to sample acquisition, conducted lab work, analyzed and interpreted the data and wrote the manuscript. JW designed the study, contributed to the analysis and interpretation of the data and reviewed and edited the manuscript. HN, MC and FK, RB, SK and KM contributed to sample acquisition and reviewed and edited the manuscript. All authors read and approved the final manuscript.

## Availability of data and materials

Sequences analyzed during the current study have been deposited to NCBI GenBank (http://www.ncbi.nlm.nih.gov/genbank) under the GenBank accession #s: PQ137681 - PQ137748.

## REFERENCES

Abduriyim S, Nishita Y, Kosintsev PA, Raichev E, Väinölä R, Kryukov AP, Abramov A V., Kaneko Y, Masuda R. 2019. Evolution of MHC class I genes in Eurasian badgers, genus Meles (Carnivora, Mustelidae). Heredity (Edinb). doi:10.1038/s41437-018-0100-3.

Abduriyim S, Zou DH, Zhao H. 2019. Origin and evolution of the major histocompatibility complex class I region in eutherian mammals. Ecol Evol. doi:10.1002/ece3.5373.

Adams EJ, Luoma AM. 2013. The adaptable major histocompatibility complex (MHC) fold: Structure and function of nonclassical and mhc class I-like molecules. Annu Rev Immunol. doi:10.1146/annurev-immunol-032712-095912.

Aguilar A, Roemer G, Debenham S, Binns M, Garcelon D, Wayne RK. 2004. High MHC diversity maintained by balancing selection in an otherwise genetically monomorphic mammal. Proc Natl Acad Sci U S A. doi:10.1073/pnas.0306582101.

Alexander KA, Laver PN, Williams MC, Sanderson CE, Kanipe C, Palmer M V. 2018. Pathology of the Emerging Mycobacterium tuberculosis Complex Pathogen, Mycobacterium mungi, in the Banded Mongoose (Mungos mungo). Vet Pathol. doi:10.1177/0300985817741730.

Alexander KA, Sanderson CE, Larsen MH, Robbe-Austerman S, Williams MC, Palmer M V. 2016. Emerging tuberculosis pathogen hijacks social communication behavior in the group-living banded mongoose (Mungos mungo). MBio. doi:10.1128/mBio.00281-16.

Alexander KA, Sanderson CE, Laver PN. 2015. Novel Mycobacterium tuberculosis complex spp. in group-living African mammals. In: Tuberculosis, leprosy and mycobacterial diseases of man and animals: the many hosts of mycobacteria.

Alfonso C, Karlsson L. 2000. Nonclassical MHC Class II Molecules. Annu Rev Immunol. doi:10.1146/annurev.immunol.18.1.113.

Aphalo P. 2024. gginnards: Explore the Innards of “ggplot2” Objects. R package version 0.2.0. https://cran.r-project.org/package=gginnards.

ASAB Ethical Committee, ABS Animal Care Committee. 2022. Guidelines for the treatment of animals in behavioural research and teaching. Anim Behav. doi:10.1016/s0003-3472(21)00389-4.

Babik W, Durka W, Radwan J. 2005. Sequence diversity of the MHC DRB gene in the Eurasian beaver (Castor fiber). Mol Ecol. doi:10.1111/j.1365-294X.2005.02751.x.

Bergström T, Gyllensten U. 1995. Evolution of Mhc Class II Polymorphism: The Rise and Fall of Class II Gene Function in Primates. Immunol Rev. doi:10.1111/j.1600-065X.1995.tb00668.x.

Bitarello BD, De Filippo C, Teixeira JC, Schmidt JM, Kleinert P, Meyer D, Andres AM. 2018. Signatures of long-term balancing selection in human genomes. Genome Biol Evol. doi:10.1093/gbe/evy054.

Bjorkman PJ, Saper MA, Samraoui B, Bennett WS, Strominger JL, Wiley DC. 1987. Structure of the human class I histocompatibility antigen, HLA-A2. Nature. doi:10.1038/329506a0.

Boni MF, Posada D, Feldman MW. 2007. An exact nonparametric method for inferring mosaic structure in sequence triplets. Genetics. doi:10.1534/genetics.106.068874.

Braud VM, Allan DS, McMichael AJ. 1999. Functions of nonclassical MHC and non-MHC-encoded class I molecules. Curr Opin Immunol. doi:10.1016/S0952-7915(99)80018-1.

Brown JH, Jardetzky TS, Gorga JC, Stern LJ, Urban RG, Strominger JL, Wiley DC. 1993. Three-dimensional structure of the human class II histocompatibility antigen HLA-DR1. Nature. doi:10.1038/364033a0.

Brüns AC, Tanner M, Williams MC, Botha L, O’Brien A, Fosgate GT, van Helden PD, Clarke J, Michel AL. 2017. Diagnosis and implications of Mycobacterium bovis infection in banded mongooses (Mungos mungo) in the Kruger National Park, South Africa. J Wildl Dis. doi:10.7589/2015-11-318.

Bürkner PC. 2017. brms: An R package for Bayesian multilevel models using Stan. J Stat Softw. doi:10.18637/jss.v080.i01.

Burnett RC, Geraghty DE. 1995. Structure and expression of a divergent canine class I gene. J Immunol. doi:10.4049/jimmunol.155.9.4278.

Cant MA, Nichols HJ, Thompson FJ, Vitikainen E. 2016. Banded mongooses: Demography, life history, and social behavior. In: Cooperative Breeding in Vertebrates: Studies of Ecology, Evolution, and Behavior.

Castro-Prieto A, Wachter B, Melzheimer J, Thalwitzer S, Sommer S. 2011. Diversity and evolutionary patterns of immune genes in free-ranging Namibian leopards (Panthera pardus pardus). In: Journal of Heredity.

Castro-Prieto A, Wachter B, Sommer S. 2011. Cheetah paradigm revisited: MHC diversity in the world’s largest free-ranging population. Mol Biol Evol. doi:10.1093/molbev/msq330.

Clarke B, Kirby DRS. 1966. Maintenance of histocompatibility polymorphisms. Nature. 211(5052):999–1000. doi:10.1038/211999a0.

Coker, O. M., Osaiyuw, O. H., Fatoki AO. 2023. Major Histocompatibility Complex (MHC) Diversity and its implications in human and wildlife health and Conservation. Genet Biodivers J. 7(2):1–11.

D’Souza MP, Adams E, Altman JD, Birnbaum ME, Boggiano C, Casorati G, Chien YH, Conley A, Eckle SBG, Früh K, et al. 2019. Casting a wider net: Immunosurveillance by nonclassical MHC molecules. PLoS Pathog. doi:10.1371/journal.ppat.1007567.

Drake GJC, Kennedy LJ, Auty HK, Ryvar R, Ollier WER, Kitchener AC, Freeman AR, Radford AD. 2004. The use of reference strand-mediated conformational analysis for the study of cheetah (Acinonyx jubatus) feline leucocyte antigen class II DRB polymorphisms. Mol Ecol. doi:10.1046/j.1365-294X.2003.02027.x.

Ebert D, Fields PD. 2020. Host–parasite co-evolution and its genomic signature. Nat Rev Genet. doi:10.1038/s41576-020-0269-1.

Faircloth BC, Glenn TC. 2012. Not all sequence tags are created equal: Designing and validating sequence identification tags robust to indels. PLoS One. doi:10.1371/journal.pone.0042543.

Gibbs MJ, Armstrong JS, Gibbs AJ. 2000. Sister-scanning: A Monte Carlo procedure for assessing signals in rebombinant sequences. Bioinformatics. doi:10.1093/bioinformatics/16.7.573.

Gutierrez-Espeleta GA, Hedrick PW, Kalinowski ST, Garrigan D, Boyce WM. 2001. Is the decline of desert bighorn sheep from infectious disease the result of low MHC variation? Heredity (Edinb). doi:10.1046/j.1365-2540.2001.00853.x.

Hedrick PW. 2003. A heterozygote advantage. Science (80-). 302(5642):57–57.

Hoang DT, Chernomor O, Von Haeseler A, Minh BQ, Vinh LS. 2018. UFBoot2: Improving the ultrafast bootstrap approximation. Mol Biol Evol. doi:10.1093/molbev/msx281.

Hodge SJ, Bell MBV, Cant MA. 2011. Reproductive competition and the evolution of extreme birth synchrony in a cooperative mammal. Biol Lett. doi:10.1098/rsbl.2010.0555.

Huang K, Zhang P, Dunn DW, Wang T, Mi R, Li B. 2019. Assigning alleles to different loci in amplifications of duplicated loci. Mol Ecol Resour. doi:10.1111/1755-0998.13036.

Hughes AL, Nei M. 1989. Nucleotide substitution at major histocompatibility complex class II loci: evidence for overdominant selection. Proc Natl Acad Sci U S A. doi:10.1073/pnas.86.3.958.

IUCN. 2024. The IUCN Red List of Threatened Species. Version 2024-1. https://www.iucnredlist.org.

Jordan NR, Mwanguhya F, Kyabulima S, Rüedi P, Cant MA. 2010. Scent marking within and between groups of wild banded mongooses. J Zool. doi:10.1111/j.1469-7998.2009.00646.x.

Kalyaanamoorthy S, Minh BQ, Wong TKF, Von Haeseler A, Jermiin LS. 2017. ModelFinder: Fast model selection for accurate phylogenetic estimates. Nat Methods. doi:10.1038/nmeth.4285.

Kamiya T, O’Dwyer K, Westerdahl H, Senior A, Nakagawa S. 2014. A quantitative review of MHC-based mating preference: The role of diversity and dissimilarity. Mol Ecol. 23(21):5151–5163. doi:10.1111/mec.12934.

Katoh K, Misawa K, Kuma KI, Miyata T. 2002. MAFFT: A novel method for rapid multiple sequence alignment based on fast Fourier transform. Nucleic Acids Res. doi:10.1093/nar/gkf436.

Katoh K, Standley DM. 2013. MAFFT multiple sequence alignment software version 7: Improvements in performance and usability. Mol Biol Evol. doi:10.1093/molbev/mst010.

Kaufman J. 2018. Unfinished Business: Evolution of the MHC and the Adaptive Immune System of Jawed Vertebrates. Annu Rev Immunol. doi:10.1146/annurev-immunol-051116-052450.

Kaufman J, Salomonsen J, Flajnik M. 1994. Evolutionary conservation of MHC class I and class II molecules - different yet the same. Semin Immunol. 6(6):411–424. doi:10.1006/smim.1994.1050.

Kelley J, Walter L, Trowsdale J. 2005. Comparative genomics of major histocompatibility complexes. Immunogenetics. doi:10.1007/s00251-004-0717-7.

Khera M, Arbuckle K, Hoffman JI, Sanderson JL, Cant MA, Nichols HJ. 2021. Cooperatively breeding banded mongooses do not avoid inbreeding through familiarity-based kin recognition. Behav Ecol Sociobiol. doi:10.1007/s00265-021-03076-3.

Klein J. 1986. Natural history of the major histocompatibility complex. New York: John Wiley & Sons.

Klein J. 1987. Origin of major histocompatibility complex polymorphism: The trans-species hypothesis. Hum Immunol. doi:10.1016/0198-8859(87)90066-8.

Knapp LA. 2005. The ABCs of MHC. Evol Anthropol. doi:10.1002/evan.20038.

Kuduk K, Babik W, Bojarska K, Śliwińska EB, Kindberg J, Taberlet P, Swenson JE, Radwan J. 2012. Evolution of major histocompatibility complex class i and class II genes in the brown bear. BMC Evol Biol. doi:10.1186/1471-2148-12-197.

Kumar S, Stecher G, Li M, Knyaz C, Tamura K. 2018. MEGA X: Molecular evolutionary genetics analysis across computing platforms. Mol Biol Evol. doi:10.1093/molbev/msy096.

Lüdecke D. 2018. ggeffects: Tidy Data Frames of Marginal Effects from Regression Models. J Open Source Softw. doi:10.21105/joss.00772.

Maccari G, Robinson J, Bontrop RE, Otting N, de Groot NG, Ho CS, Ballingall KT, Marsh SGE, Hammond JA. 2018. IPD-MHC: nomenclature requirements for the non-human major histocompatibility complex in the next-generation sequencing era. Immunogenetics. doi:10.1007/s00251-018-1072-4.

Marshall HH, Johnstone RA, Thompson FJ, Nichols HJ, Wells D, Hoffman JI, Kalema-Zikusoka G, Sanderson JL, Vitikainen EIK, Blount JD, et al. 2021. A veil of ignorance can promote fairness in a mammal society. Nat Commun. doi:10.1038/s41467-021-23910-6.

Martin D, Rybicki E. 2000. RDP: Detection of recombination amongst aligned sequences. Bioinformatics. doi:10.1093/bioinformatics/16.6.562.

Martin DP, Murrell B, Golden M, Khoosal A, Muhire B. 2015. RDP4: Detection and analysis of recombination patterns in virus genomes. Virus Evol. doi:10.1093/ve/vev003.

Martin DP, Murrell B, Khoosal A, Muhire B. 2017. Detecting and analyzing genetic recombination using RDP4. In: Methods in Molecular Biology.

Murrell B, Moola S, Mabona A, Weighill T, Sheward D, Kosakovsky Pond SL, Scheffler K. 2013. FUBAR: A fast, unconstrained bayesian AppRoximation for inferring selection. Mol Biol Evol. doi:10.1093/molbev/mst030.

Murrell B, Wertheim JO, Moola S, Weighill T, Scheffler K, Kosakovsky Pond SL. 2012. Detecting individual sites subject to episodic diversifying selection. PLoS Genet. doi:10.1371/journal.pgen.1002764.

Neefjes J, Jongsma MLM, Paul P, Bakke O. 2011. Towards a systems understanding of MHC class I and MHC class II antigen presentation. Nat Rev Immunol. 11(12):823–836. doi:10.1038/nri3084.

Nei M, Gojobori T. 1986. Simple methods for estimating the numbers of synonymous and nonsynonymous nucleotide substitutions. Mol Biol Evol.

Nei M, Gu X, Sitnikova T. 1997. Evolution by the birth-and-death process in multigene families of the vertebrate immune system. Proc Natl Acad Sci U S A. doi:10.1073/pnas.94.15.7799.

Nguyen LT, Schmidt HA, Von Haeseler A, Minh BQ. 2015. IQ-TREE: A fast and effective stochastic algorithm for estimating maximum-likelihood phylogenies. Mol Biol Evol. doi:10.1093/molbev/msu300.

Nichols HJ, Cant MA, Hoffman JI, Sanderson JL. 2014. Evidence for frequent incest in a cooperatively breeding mammal. Biol Lett. doi:10.1098/rsbl.2014.0898.

Nichols HJ, Cant MA, Sanderson JL. 2015. Adjustment of costly extra-group paternity according to inbreeding risk in a cooperative mammal. Behav Ecol. doi:10.1093/beheco/arv095.

Okamura K, Dijkstra JM, Tsukamoto K, Grimholt U, Wiegertjes GF, Kondow A, Yamaguchi H, Hashimoto K. 2021. Discovery of an ancient MHC category with both class I and class II features. Proc Natl Acad Sci U S A. doi:10.1073/pnas.2108104118.

Okano M, Miyamae J, Suzuki S, Nishiya K, Katakura F, Kulski JK, Moritomo T, Shiina T. 2020. Identification of Novel Alleles and Structural Haplotypes of Major Histocompatibility Complex Class I and DRB Genes in Domestic Cat (Felis catus) by a Newly Developed NGS-Based Genotyping Method. Front Genet. doi:10.3389/fgene.2020.00750.

Padidam M, Sawyer S, Fauquet CM. 1999. Possible emergence of new geminiviruses by frequent recombination. Virology. doi:10.1006/viro.1999.0056.

Pierini F, Lenz TL. 2018. Divergent allele advantage at human MHC genes: Signatures of past and ongoing selection. Mol Biol Evol. doi:10.1093/molbev/msy116.

Piertney SB, Oliver MK. 2006. The evolutionary ecology of the major histocompatibility complex. Heredity (Edinb). doi:10.1038/sj.hdy.6800724.

Pokorny I, Sharma R, Goyal SP, Mishra S, Tiedemann R. 2010. MHC class I and MHC class II DRB gene variability in wild and captive Bengal tigers (Panthera tigris tigris). Immunogenetics. doi:10.1007/s00251-010-0475-7.

Posada D, Crandall KA. 2001. Evaluation of methods for detecting recombination from DNA sequences: Computer simulations. Proc Natl Acad Sci U S A. doi:10.1073/pnas.241370698.

R Core Team. 2023. R Core Team 2023 R: A language and environment for statistical computing. R foundation for statistical computing. https://www.R-project.org/. R Found Stat Comput.

Radwan J, Babik W, Kaufman J, Lenz TL, Winternitz J. 2020. Advances in the Evolutionary Understanding of MHC Polymorphism. Trends Genet. doi:10.1016/j.tig.2020.01.008.

Radwan J, Biedrzycka A, Babik W. 2010. Does reduced MHC diversity decrease viability of vertebrate populations? Biol Conserv. doi:10.1016/j.biocon.2009.07.026.

Rammensee HG, Bachmann J, Stevanović S. 2013. MHC ligands and peptide motifs. Springer Science & Business Media.

Rao X, Hoof I, Fontaine Costa AICA, Van Baarle D, Keşmir C. 2011. HLA class I allele promiscuity revisited. Immunogenetics. 63(11):691–701. doi:10.1007/s00251-011-0552-6.

Reche PA, Reinherz EL. 2003. Sequence variability analysis of human class I and class II MHC molecules: Functional and structural correlates of amino acid polymorphisms. J Mol Biol. doi:10.1016/S0022-2836(03)00750-2.

Reche PA, Reinherz EL. 2007. Definition of MHC supertypes through clustering of MHC peptide-binding repertoires. Methods Mol Biol. doi:10.1007/978-1-60327-118-9_11.

Revell LJ. 2012. phytools: An R package for phylogenetic comparative biology (and other things). Methods Ecol Evol. doi:10.1111/j.2041-210X.2011.00169.x.

Revell LJ. 2024. phytools 2.0: an updated R ecosystem for phylogenetic comparative methods (and other things). PeerJ. doi:10.7717/peerj.16505.

Rozas J, Ferrer-Mata A, Sanchez-DelBarrio JC, Guirao-Rico S, Librado P, Ramos-Onsins SE, Sanchez-Gracia A. 2017. DnaSP 6: DNA sequence polymorphism analysis of large data sets. Mol Biol Evol. doi:10.1093/molbev/msx248.

Salminen MO, Carr JK, Burke DS, Mccutchan FE. 1995. Identification of Breakpoints in Intergenotypic Recombinants of HIV Type 1 by Bootscanning. AIDS Res Hum Retroviruses. doi:10.1089/aid.1995.11.1423.

Sanderson JL, Wang J, Vitikainen EIK, Cant MA, Nichols HJ. 2015. Banded mongooses avoid inbreeding when mating with members of the same natal group. Mol Ecol. doi:10.1111/mec.13253.

Saper MA, Bjorkman PJ, Wiley DC. 1991. Refined structure of the human histocompatibility antigen HLA-A2 at 2.6 Å resolution. J Mol Biol. doi:10.1016/0022-2836(91)90567-P.

Satta Y, O’Huigin C, Takahata N, Klein J. 1994. Intensity of natural selection at the major histocompatibility complex loci. Proc Natl Acad Sci U S A. doi:10.1073/pnas.91.15.7184.

Sebastian A, Herdegen M, Migalska M, Radwan J. 2016. Amplisas: A web server for multilocus genotyping using next-generation amplicon sequencing data. Mol Ecol Resour. doi:10.1111/1755-0998.12453.

Singh PB. 1998. The present status of the “carrier hypothesis” for chemosensory recognition of genetic individuality. Genetica. doi:10.1023/A:1026475118901.

Smith JM. 1992. Analyzing the mosaic structure of genes. J Mol Evol. doi:10.1007/BF00182389.

Sommer S. 2005. The importance of immune gene variability (MHC) in evolutionary ecology and conservation. Front Zool. doi:10.1186/1742-9994-2-16.

Takahashi K, Rooney AP, Nei M. 2000. Origins and divergence times of mammalian class II MHC gene clusters. In: Journal of Heredity.

Takahata N, Nei M. 1990. Allelic genealogy under overdominant and frequency-dependent selection and polymorphism of major histocompatibility complex loci. Genetics. doi:10.1093/genetics/124.4.967.

Těšický M, Vinkler M. 2015. Trans-Species Polymorphism in Immune Genes: General Pattern or MHC-Restricted Phenomenon? J Immunol Res. doi:10.1155/2015/838035.

Trifinopoulos J, Nguyen LT, von Haeseler A, Minh BQ. 2016. W-IQ-TREE: a fast online phylogenetic tool for maximum likelihood analysis. Nucleic Acids Res. doi:10.1093/NAR/GKW256.

Ujvari B, Belov K. 2011. Major histocompatibility complex (MHC) markers in conservation biology. Int J Mol Sci. doi:10.3390/ijms12085168.

Upham NS, Esselstyn JA, Jetz W. 2019. Inferring the mammal tree: Species-level sets of phylogenies for questions in ecology, evolution, and conservation. PLoS Biol. doi:10.1371/journal.pbio.3000494.

Weaver S, Shank SD, Spielman SJ, Li M, Muse S V., Kosakovsky Pond SL. 2018. Datamonkey 2.0: A modern web application for characterizing selective and other evolutionary processes. Mol Biol Evol. doi:10.1093/molbev/msx335.

Wells DA, Cant MA, Hoffman JI, Nichols HJ. 2020. Inbreeding depresses altruism in a cooperative society. Ecol Lett. doi:10.1111/ele.13578.

Wells DA, Cant MA, Nichols HJ, Hoffman JI. 2018. A high-quality pedigree and genetic markers both reveal inbreeding depression for quality but not survival in a cooperative mammal. Mol Ecol. doi:10.1111/mec.14570.

Wickham H. 2016. ggplot2: Elegant Graphics for Data Analysis. Second Edition. Springer.

Winternitz J, Abbate JL, Huchard E, Havlíček J, Garamszegi LZ. 2017. Patterns of MHC-dependent mate selection in humans and nonhuman primates: a meta-analysis. Mol Ecol. 26(2):668–688.

Winternitz J, Chakarov N, Rinaud T, Ottensmann M, Krüger O. 2023. High functional allelic diversity and copy number in both MHC classes in the common buzzard. BMC Ecol Evol. doi:10.1186/s12862-023-02135-9.

Winternitz JC, Abbate JL. 2022. The genes of attraction: Mating behavior, immunogenetic variation, and parasite resistance. In: Animal Behavior and Parasitism.

Yu G, Smith DK, Zhu H, Guan Y, Lam TTY. 2017. ggtree: an r package for visualization and annotation of phylogenetic trees with their covariates and other associated data. Methods Ecol Evol. doi:10.1111/2041-210X.12628.

Yuhki N, Heidecker GF, O’Brien SJ. 1989. Characterization of MHC cDNA clones in the domestic cat. Diversity and evolution of class I genes. J Immunol. doi:10.4049/jimmunol.142.10.3676.

Yuhki N, Mullikin JC, Beck T, Stephens R, O’Brien SJ. 2008. Sequences, annotation and single nucleotide polymorphism of the major histocompatibility complex in the domestic cat. PLoS One. doi:10.1371/journal.pone.0002674.

Yuhki N, O’Brien SJ. 1990. DNA variation of the mammalian major histocompatibility complex reflects genomic diversity and population history. Proc Natl Acad Sci U S A. doi:10.1073/pnas.87.2.836.

Yuhki N, O’Brien SJ. 1994. Exchanges of short polymorphic DNA segments predating speciation in feline major histocompatibility complex class I genes. J Mol Evol. doi:10.1007/BF00178246.

Yuhki N, O’Brien SJ. 1997. Nature and origin of polymorphism in feline MHC class II DRA and DRB genes. J Immunol. doi:10.4049/jimmunol.158.6.2822.

Zhang C, Anderson A, DeLisi C. 1998. Structural principles that govern the peptide-binding motifs of class I MHC molecules. J Mol Biol. doi:10.1006/jmbi.1998.1982.

Zhang Z, Schwartz S, Wagner L, Miller W. 2000. A greedy algorithm for aligning DNA sequences. J Comput Biol. doi:10.1089/10665270050081478.

Zhu Y, Sun DD, Ge YF, Yu B, Chen YY, Wan QH. 2013. Isolation and characterization of class I MHC genes in the giant panda (Ailuropoda melanoleuca). Chinese Sci Bull. doi:10.1007/s11434-012-5582-4.

Zinkernagel RM, Doherty PC. 1975. H 2 compatibility requirement for T cell mediated lysis of target cells infected with lymphocytic choriomeningitis virus. Different cytotoxic T cell specificities are associated with structures coded for in H 2K or H 2D. J Exp Med. doi:10.1084/jem.141.6.1427.

