## Supplementary figures and images for "Characterization of both major histocompatibility complex classes in a wild social mammal: the banded mongoose"

### Figure S1

#### Flow chart – artefact identification

blue = keep sequence, red = remove sequence

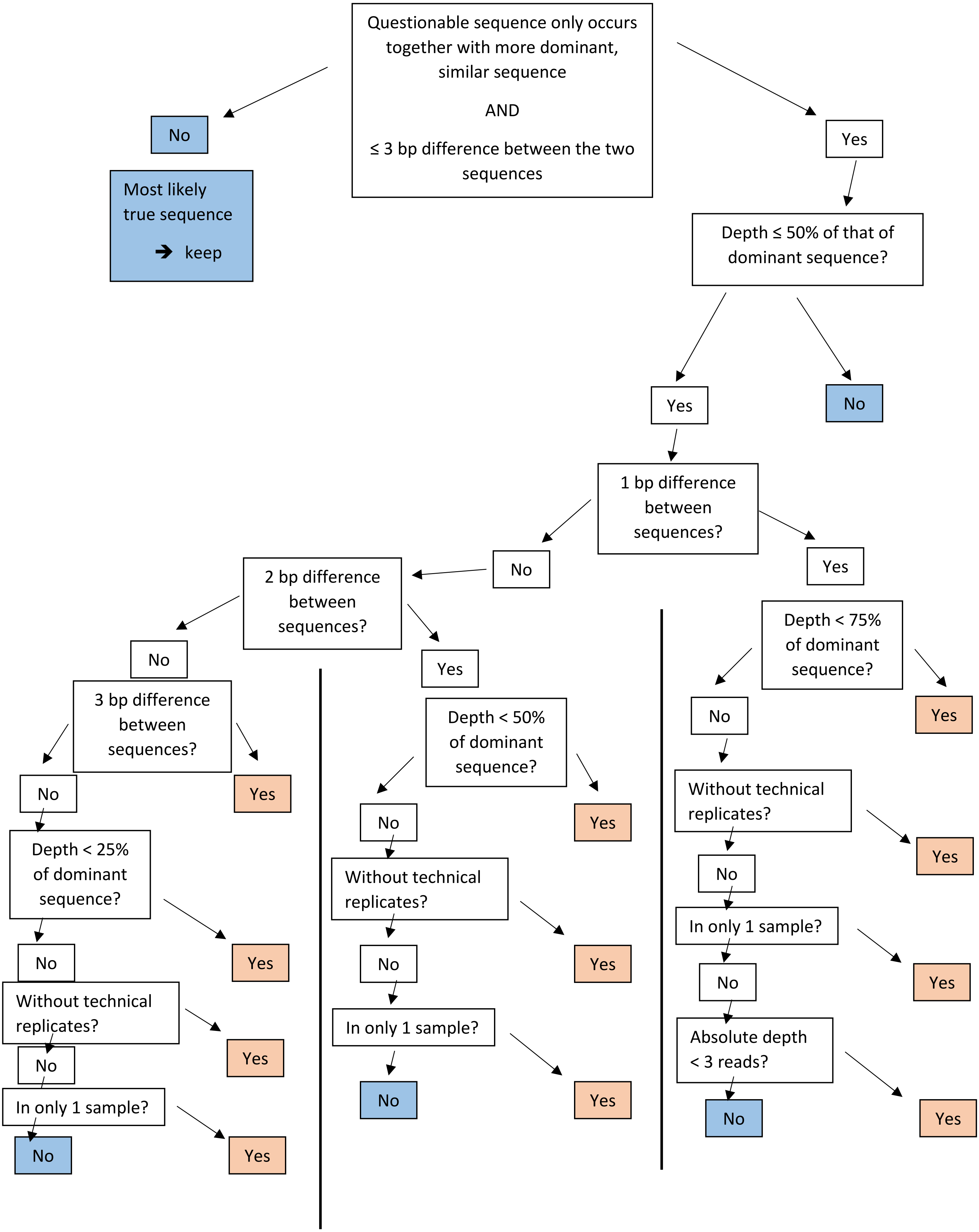
