## Supplementary material for "Characterization of both major histocompatibility complex classes in a wild social mammal: the banded mongoose": Figure S2

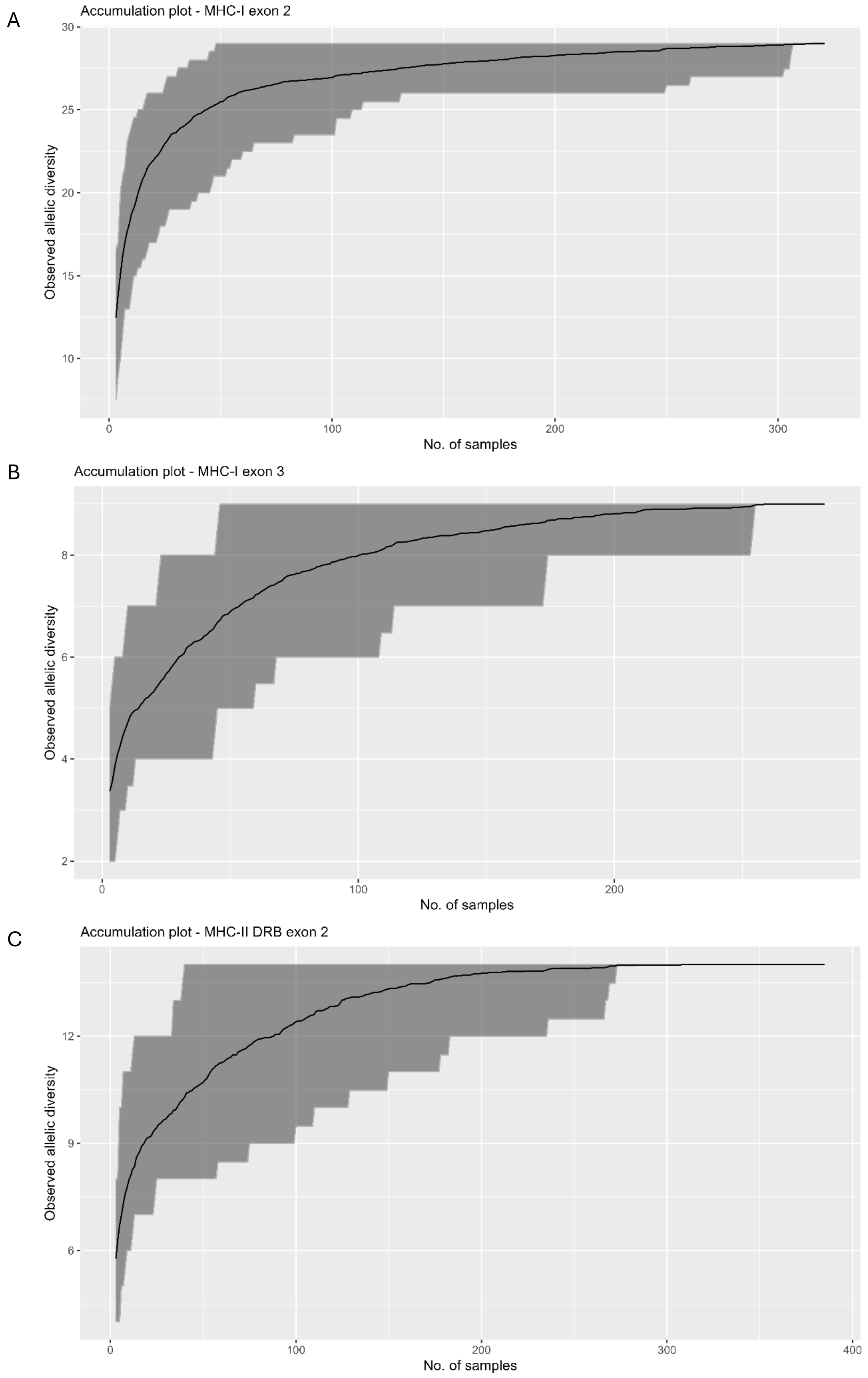


**Figure S2** Accumulation curves. Displayed are the accumulation curves for the different MHC classes and exons. The analysis shows the estimated recovered MHC diversity per sample site for MHC-I exon 2

**(a)**, for MHC-I exon 3 **(b)**, and for MHC-II DRB exon 2 **(c)**.
