## Supplementary material for "Characterization of both major histocompatibility complex classes in a wild social mammal: the banded mongoose": Figure S3

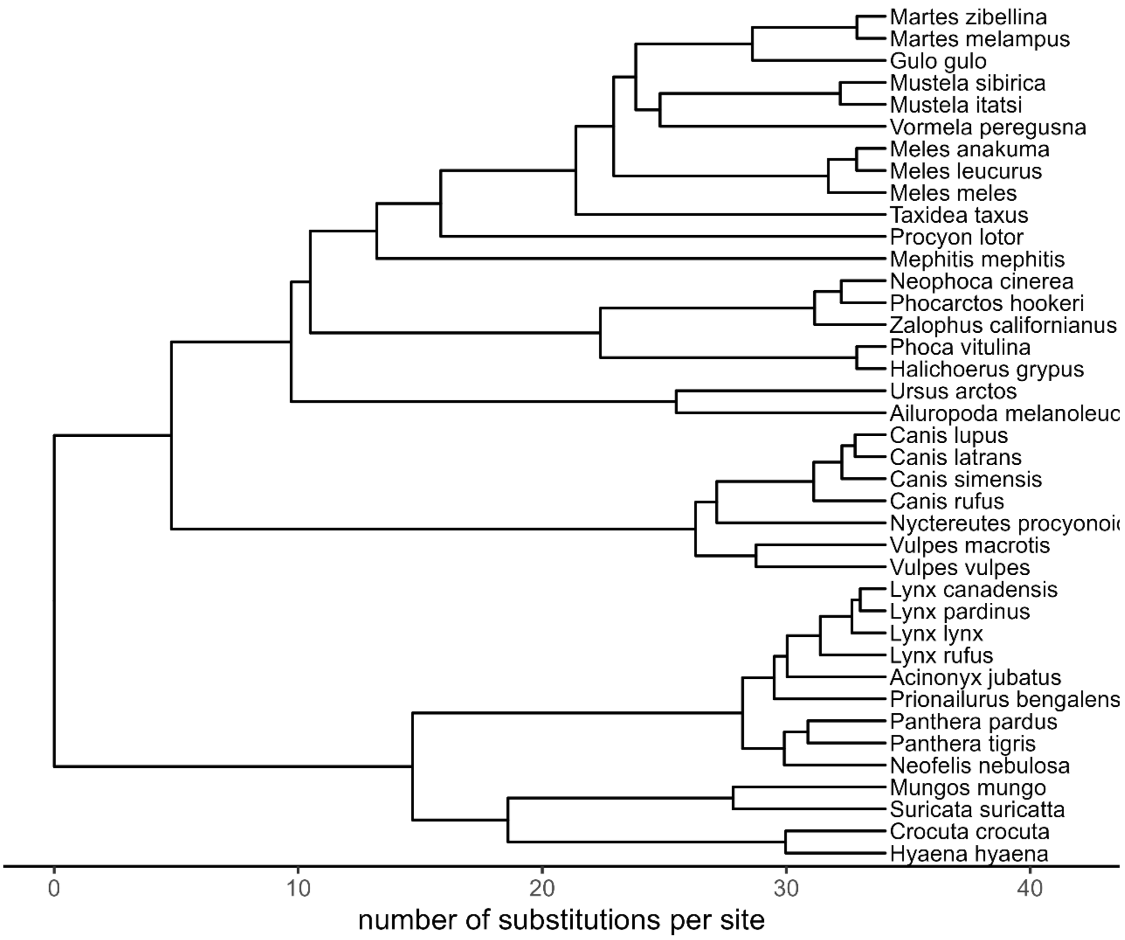


**Figure S3 Consensus phylogenetic tree** Phylogenetic tree used for comparison of MHC diversity for the species listed.
