## Supplementary Methods for "Characterization of both major histocompatibility complex classes in a wild social mammal: the banded mongoose"

**Assigning alleles to loci**

We used MHC typer V1.1, which uses a maximum-likelihood approach for reconstructing haplotypes while considering null alleles or copy number variation (CNV), identical alleles shared between loci as well as deviations from Hardy-Weinberg-Equilibrium (Huang et al. 2019). The protocol used was the recommended 2-step procedure (Huang et al. 2019): step 1: random initial solution for #loci=8 for MHC-I exon 2, 3 for MHC-I exon 3 and 4 for MHC-II DRB exon 2, number of repeats=10, chain length=500, initial temperature=0.01, final temperature=0.00001, anneal coefficient=0.99, max iteration=30, min freq. diff=0.002, penalty of missing=-2, penalty of mismatch=-80, taboo coef.=1; Step 2: initial solution for loci=solution from best run of Step 1, number of repeats=300, initial temperature=0.0001, final temperature=0.000001, consider null alleles, consider deviation from HWE, initial null allele freq.=0.05, penalty of missing=0, penalty of mismatch=-1000, taboo coef.=1.00001. We considered an assignment to loci as reliable when a run resulted in the same assignment at least twice with an optimal Bayesian information criterion (BIC).

Huang K, Zhang P, Dunn DW, Wang T, Mi R, Li B. 2019. Assigning alleles to different loci in amplifications of duplicated loci. Mol Ecol Resour. doi:10.1111/1755-0998.13036.
